# Conversion of silent synapses to AMPA receptor-mediated functional synapses in human cortical organoids

**DOI:** 10.1101/2024.12.21.629885

**Authors:** Masatoshi Nishimura, Tomoki Kodera, Shota Adachi, Akinori Y. Sato, Ryosuke F. Takeuchi, Hiroshi Nonaka, Itaru Hamachi, Fumitaka Osakada

**Author notes:** Correspondence: Fumitaka Osakada, Ph.D., Laboratory of Cellular Pharmacology, Graduate School of Pharmaceutical Sciences, Nagoya, University, Nagoya, Japan.

## Abstract

Despite the crucial role of synaptic connections and neural activity in the development and organization of cortical circuits, the mechanisms underlying the formation of functional synaptic connections in the developing human cerebral cortex remain unclear. We investigated the development of α-amino-3-hydroxy-5-methyl-4-isoxazolepropionic acid receptor (AMPAR)-mediated synaptic transmission using human cortical organoids (hCOs) derived from induced pluripotent stem cells. Two-photon Ca^2^⁺ imaging revealed an increase in the frequency and amplitude of spontaneous activity in hCOs on day 80 compared to day 50. Additionally, spontaneous neural activity in late-stage hCOs, but not in early-stage hCOs, was blocked by N-methyl-D-aspartate receptor (NMDAR) and AMPAR antagonists. However, transsynaptic circuit tracing with G-deleted rabies viral vectors indicated a similar number of synaptic connections in early- and late-stage hCOs. Notably, chemical labeling demonstrated a significant increase in AMPAR expression on the postsynaptic membrane and colocalization with NMDAR in late-stage hCOs. These results suggest that hCOs progressively organize excitatory synaptic transmission, concurrent with the transition from silent synapses lacking AMPARs to functional synapses containing NMDARs and AMPARs. This *in vitro* model of human cortical circuits derived from induced pluripotent stem cells reflects the developmental programs underlying physiological transitions, providing valuable insights into human corticogenesis and neurodevelopmental disorders.

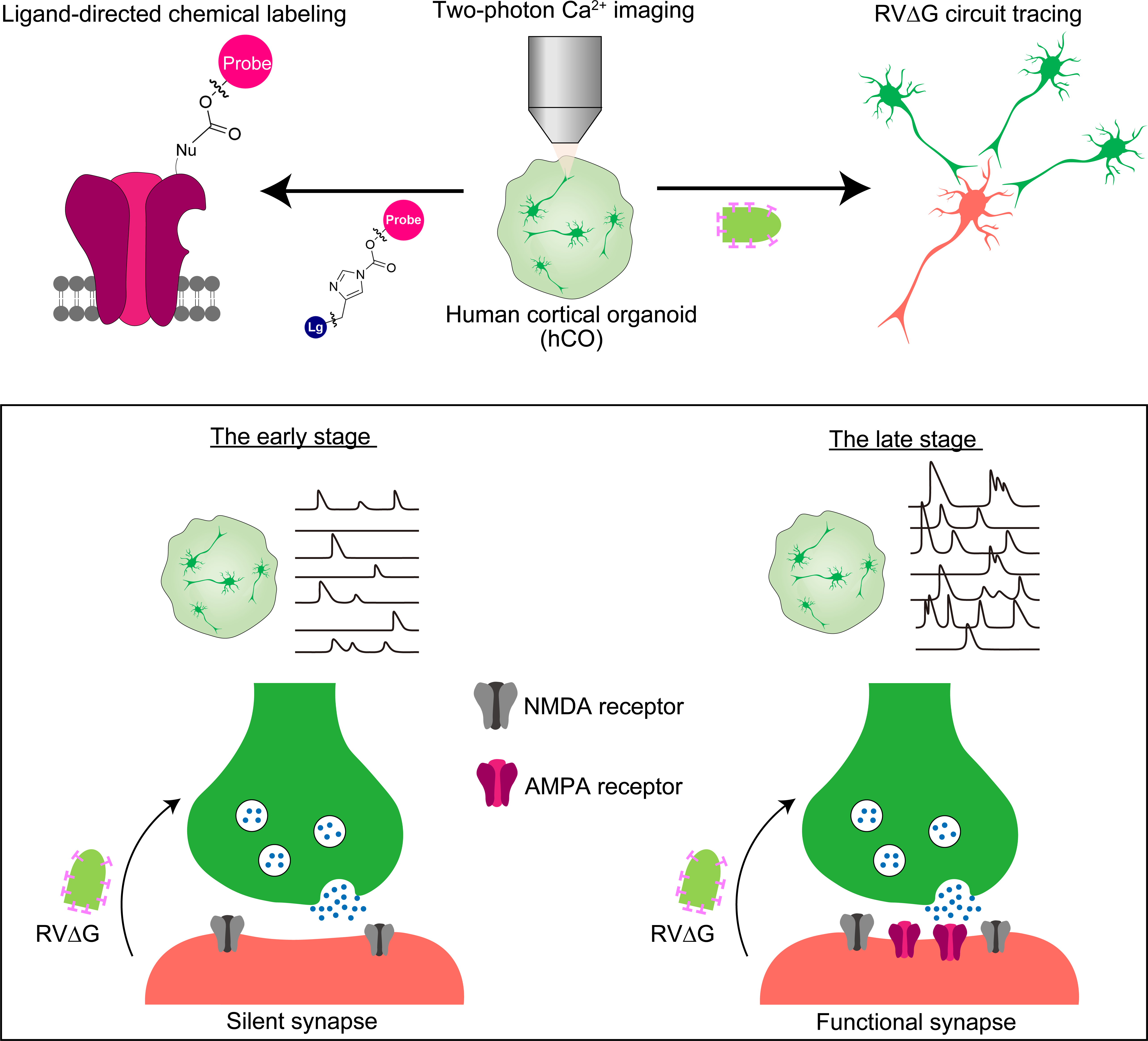

## Introduction

The cerebral cortex, the largest part of the human brain, governs higher-order functions, such as sensory perception, motor control, and cognition (Kolk and Rakic, 2022; Van Essen et al., 2019). These functions arise from neural circuits formed during early development when neural progenitors located in the ventricular zone radially migrate and differentiate into cortical neurons to form the cortical plate. Subsequent spontaneous neural activity regulates neuronal survival, axonal growth, and synapse formation (Riva et al., 2019; Spitzer, 2006; Tagawa and Hirano, 2012) and is transmitted through gap junctions and synapses (Pereda, 2014). Initially, new synapses in the developing brain are silent, featuring the expression of only N-methyl-D-aspartate receptors (NMDARs). However, synapses mature into functionally active synapses with NMDARs and α-amino-3-hydroxy-5-methyl-4-isoxazolepropionic acid receptors (AMPARs), enabling functional synaptic transmission (Hanse et al., 2013; Kanold et al., 2019). Neural activity shifts from asynchronous to synchronous, refining neural circuits in the developing cortex (Molnár et al., 2020). These activity changes contribute to primitive patterning that is responsible for processing different sensory modalities in the cerebral cortex (Bragg-Gonzalo et al., 2021; Wu et al., 2024). Although rodent models have been pivotal in elucidating these aspects of cortical development, they cannot fully capture the unique features of the human cerebral cortex. The human cerebral cortex has unique structures and exhibits vulnerabilities to neurological and psychiatric disorders, including autism and schizophrenia (Lancaster, 2024). Therefore, to understand human corticogenesis and neurodevelopmental disorders, it is necessary to create models that accurately replicate human cortical development.

Recent advances in neural organoid technologies have enabled the modeling of human brain development *in vitro* (Lancaster et al., 2013; Pașca, 2018; Sasai, 2013). Generating cortical organoids from pluripotent stem cells has facilitated the investigation of key aspects of cerebral cortex development, including cell type diversity, cytoarchitecture, neural activity, and synaptic formation (Eiraku et al., 2008; Paşca et al., 2015; Velasco et al., 2019). However, how neural activity and synaptic connections are organized during human corticogenesis remains unclear.

In this study, we generated cortical organoids (hCOs) from human induced pluripotent stem cells (hiPSCs) and employed a combination of two-photon Ca^2+^ imaging, transsynaptic viral tracing (Osakada et al., 2011; Wickersham et al., 2007), and ligand-directed chemical labeling (Nonaka et al., 2024; Wakayama et al., 2017) to investigate developmental changes in neural activity and synaptic connections. We quantitatively assessed synaptic connections through G-deleted rabies viral vectors (RVΔGs), adeno-associated vectors (AAVs), and cell sorting technologies. A chemical labeling method allowed us to analyze NMDARs and AMPARs on the cell surface in hCOs. Our findings revealed AMPAR-mediated synaptic transmission driven by the transition of synapses from silent to functional states, indicating the physiological maturation of excitatory synapses in human cortical circuits in organoids. We provide an *in vitro* model of human cortical circuits that reflects the developmental programs underlying physiological transitions in synapses and neural activity. By bridging the gap with rodent models, this study contributes to our understanding of the development and pathophysiology of the human cerebral cortex.

## Materials and methods

All experiments were approved by Nagoya University and conducted in accordance with the Guidelines of Nagoya University.

### hiPSC culture

hiPSC lines (clone: 1383D6) were provided by Drs. Masato Nakagawa and Shinya Yamanaka of the Center for iPS Cell Research, Kyoto University. Human iPS cells were maintained on iMatrix 511 (Nippi Inc., Tokyo, Japan)-coated culture dishes in StemFit AK02N (Ajinomoto, Tokyo, Japan) as described previously (Ye et al., 2020). For passaging, when hiPSCs reached sub-confluency, they were treated with Accutase (Nacalai Tesque, Kyoto, Japan), dissociated into single cells, and replated at a density of 1.0 × 10^3^ cells/cm^2^ onto iMatrix 511-coated dishes in AK02N in the presence of 10 μM Y-27632 (ROCK inhibitor, Wako, Osaka, Japan) for 24 h. The medium was changed to a fresh one without Y-27632 one day after plating and every day thereafter. hiPSCs were maintained in a humidified atmosphere of 5% CO_2_ at 37°C.

### hCO generation from hiPSCs

hCOs were generated as previously described (Xiang et al., 2017) with some modifications. Briefly, single-cell suspensions of undifferentiated hiPSCs were plated into a low-adherent 96-well V-bottom plate (10,000 cells/well) in Dulbecco’s Modified Eagle’s Medium (DMEM)/Ham’s F12 + KnockOut Serum Replacement (KSR) medium (DMEM/F12, 15% KSR, 1% Minimum Essential Medium Non-Essential Amino Acid [MEM-NEAA]; 1% L-glutamine, and 100 µM β-mercaptoethanol) supplemented with 100 nM LDN-193189 (ALK2/3 inhibitor), 500 nM A-83-01 (ALK4/5/7 inhibitor), 2 µM IWR-1 *endo* (Wnt/β-catenin signal inhibitor), and 50 µM Y27632 (ROCK inhibitor). Half of the medium was replaced with fresh medium every other day. Y27632 was removed on day 4. On day 10, the organoids were transferred to spinning culture (70 rpm) in a low-adherent 6-cm dish and maintained in DMEM/F12 + Neurobasal medium (1:1 mixture of DMEM/F12 and Neurobasal medium, 0.5% N2 supplement, 1% B27 supplement minus vitamin A, 0.5% MEM-NEAA, 1% L-glutamine, 0.025% insulin, 50 µM β-mercaptoethanol, and 1% penicillin/streptomycin) supplemented with 20 ng/mL basic fibroblast growth factor (bFGF). The medium was changed every other day until day 18. From day 18 onward, organoids were maintained in DMEM/F12 + Neurobasal medium supplemented with 20 ng/mL brain-derived neurotrophic factor and 200 µM ascorbic acid, changing the medium every other day thereafter.

### Quantitative reverse-transcription polymerase chain reaction (qPCR)

Gene expression levels were evaluated using qPCR as described previously (Kodera et al., 2023; Ye et al., 2020). Total RNA was isolated using the FavorPrep Tissue Total RNA Purification Mini Kit. Subsequently, 500 ng of total RNA was reverse-transcribed using the PrimeScript RT Master Mix (Takara). The synthesized cDNA was analyzed by qPCR using the TB Green Fast qPCR Mix (Takara) on a LightCycler system (Roche). The primers used in this study are listed in the Supplementary Information.

### Sectioning and immunostaining

Organoids were fixed with 4% paraformaldehyde (PFA) in phosphate-buffered saline (PBS) at 4°C overnight and washed three times with PBS at room temperature. Fixed organoids were then transferred into 15% sucrose in PBS for two overnight incubations, followed by 30% sucrose in PBS, as described previously (Kodera et al., 2023). Organoids were transferred to cryomolds (Sakura Finetek) and embedded in a 2:1 mixture of optimal cutting temperature compound (Sakura Finetek) and a 30% sucrose solution. Organoids were cut into 20–40-µm sections using a cryostat (CM3050S, Leica). Immunostaining was performed as previously described (Kodera et al., 2023; Osakada et al., 2008). Organoid slices were reacted with Blocking One (Nacalai Tesque) for 1 h at room temperature before incubation with primary antibodies overnight at 4°C. Sections were washed with PBS three times and incubated with secondary antibodies for 90 min at room temperature. Cell nuclei were counterstained with 4′,6-diamidino-2-phenylindole (DAPI; 1.25 µg/mL). Specimens were imaged using a confocal microscope (LSM800, Zeiss) and a fluorescence macroscope (Thunder, Leica). The antibodies used, along with their working dilutions, are described in the Supplementary Information.

### Two-photon Ca^2+^ imaging

Two-photon Ca^2+^ imaging was performed as previously described (Kodera et al., 2023). The lower half of the organoids was embedded in 4% low-melting-point agarose filled with oxygenated artificial cerebrospinal fluid (ACSF; 125 mM NaCl, 2.5 mM KCl, 1.25 mM NaH_2_PO_4_•H_2_O, 25 mM NaHCO_3_, 25 mM D-glucose, 2 mM CaCl_2_, and 1 mM MgCl_2_). Imaging was conducted using a two-photon fluorescence microscope equipped with a GaAsP-type non-descanned detector and resonant scanner (A1R-MP+, Nikon) (Masaki et al., 2022). A 920-nm pulsed laser (InSight DeepSee+, Spectra-Physics) was used to excite jGCaMP7f. Time-lapse images were acquired using a 25× water immersion lens (NA: 1.1, Nikon) at a resolution of 512 × 512 pixels at 30.0 Hz. For the pharmacological approach, 1 μM tetrodotoxin (TTX) or 50 µM CNQX and 50 µM MK-801 were added to the ACSF during imaging and washed three times with ACSF.

MATLAB (MathWorks, R2022b) was used for image processing and statistical quantification. In the first step of image processing for the time-lapse data, cross-correlation– based rigid image registration (Guizar-Sicairos et al., 2008) was performed to correct for image displacement caused by motion artifacts. Changes in fluorescence intensity in individual cells were quantified. The regions of interest in individual cells were semi-manually defined using the “Cell Magic Wand” plugin in ImageJ (Scholl et al., 2017). To reduce high-frequency artifacts caused by electrical noise, the fluorescence data of jGCaMP7f were smoothed using Savitzky–Golay filters (filter width = 301 frames). The normalized fluorescence signal change (ΔF/F_0_) was computed to evaluate the activity of each neuron. The baseline of each cell (F_0_) was calculated using a low-pass percentile filter (10th percentile; cutoff frequency = 1/15 Hz). Active cells were determined based on three criteria calculated from ΔF/F_0_: (1) skewness of the ΔF/F_0_ distribution during recording > 1.0; (2) SD of ΔF/F_0_ > 0.04; and (3) ≥ 1 detected Ca^2+^ event during the recording period. A Ca^2+^ event in jGCaMP7f was defined as the peak with “MinPeakHeight = 0.2,” “MinPeakDistance = 30,” and “MinPeakWidth = 15” using the MATLAB findpeaks function. The maximum amplitude was defined as the maximum height of the Ca^2+^ event during the recording period. The correlation between the activities of recorded cells in hCOs was estimated by computing the Pearson’s correlation coefficient (r) of ΔF/F_0_ in each cell during the recording period. The mean correlation was calculated as the average correlation coefficient of all active cell pairs for each imaging site. Principal component analysis (PCA) was performed to reduce dimensionality of imaging data on days 50 and 80 for imaging sites with ≥ 10 active cells.

### AAV production

AAV vectors were generated in HEK293T cells as previously described (Suzuki et al., 2019). Briefly, AAVDJ was produced by transfecting HEK293T cells with pHelper, the AAVDJ rep/cap vector, and the genomic vector pAAV-CaMKIIa-jGCaMP7f, pAAV-hSyn-TVAmCherry, or pAAV-hSyn-TVAmCherry-P2A-oG. Transfected cells were harvested 3 days after transfection and lysed via freeze-and-thaw cycles for purification. After centrifugation, the supernatant was loaded onto gradients (15%, 25%, 40%, and 58%) of iodixanol OptiPrep (Serumwerk Bernburg). After centrifugation at 16,000 ×*g* at 4°C for 4 h, 200 µL of the 40% iodixanol fraction was collected and used for infection experiments. AAV genomic titers were quantified by qPCR. The titers of AAVDJ-CaMKIIa-jGCaMP7f, AAVDJ-hSyn-TVAmCherry, and AAVDJ-hSyn-TVAmCherry-P2A-oG were 0.314–2.17 × 10^12^, 4.07–5.45 × 10^12^, and 1.19–2.88 × 10^12^ viral genomes/mL, respectively. Virus aliquots were stored at −80°C until use.

### RVΔG production

EnvA-pseudotyped RVΔGs were produced as described previously (Okigawa et al., 2021; Osakada et al., 2011; Suzuki et al., 2019). Briefly, RVΔGs were recovered by transfecting B7GG cells with pcDNA-SAD-B19N, pcDNA-SAD-B19P, pcDNA-SAD-B19L, pcDNA-SAD-B19G, and the rabies viral genome vector pSAD-B19ΔG-EGFP using Lipofectamine 2000 (Thermo Fisher Scientific). For pseudotyping RVΔG with EnvA, BHK-EnvA cells were infected with unpseudotyped RVΔG. The virus-containing medium was concentrated through two rounds of ultracentrifugation (Optima XE-90, Beckman Coulter). The infectious titers in HEK-TVA cells were then determined. HEK293T cells were used to check for contamination with unpseudotyped RVΔG. Virus aliquots were stored at −80°C until use. The EnvA-SAD-B19ΔG-EGFP titer was 1.0–7.6 × 10^10^ infectious units/mL.

### Neural circuit tracing with RVΔG

For retrograde transsynaptic tracing in hCOs, hCOs were first infected with AAVDJ-hSyn-TVAmCherry or AAVDJ-hSyn-TVAmCherry-P2A-oG. Ten days after AAV infection, the hCOs were then infected with EnvA-RVΔG-EGFP. Another ten days after rabies viral delivery, hCOs were fixed overnight in 4% PFA in PBS at 4°C and subjected to immunostaining. For quantification, hCOs were dissociated into single cells; GFP^+^mCherry⁻ presynaptic cells and GFP^+^mCherry⁺ postsynaptic cells were counted using a cell sorter (CytoFLEX SRT, Beckman Coulter). Dead cells were excluded by gating on the forward and side scatter.

### Organoid dissociation

hCOs were enzymatically dissociated as previously described (Melliou and Diamandis, 2022) with some modifications. Briefly, hCOs were collected, washed once with PBS, and incubated with TrypLE Select (Thermo Fisher Scientific) diluted 2-fold with a 1 mM EDTA solution for 30 min at 37°C. hCOs were then dissected into small aggregates by pipetting and incubated for 15 min at 37°C to complete dissociation. After stopping the enzymatic reaction with fluorescence-activated cell sorting (FACS) buffer (PBS containing 2% fetal bovine serum), dissociated cells were centrifuged at 200 ×*g* for 5 min. The cell pellet was resuspended in the FACS buffer and filtered through a 40-µm cell strainer. The prepared sample was immediately subjected to cell sorting (CytoFLEX SRT, Beckman Coulter).

### Chemical labeling of AMPARs and NMDARs

For ligand-directed chemical AMPAR labeling (Nonaka et al., 2024; Wakayama et al., 2017), hCOs were treated with CAM2-Ax647 in DMEM/F12 + Neurobasal medium for 1 h in a humidified atmosphere of 5% CO₂ at 37°C. No additives were included in this medium to prevent labeling inhibition. To assess CAM2-Ax647 specificity, 20 µM CNQX, a competitive AMPAR antagonist, was incubated with CAM2-Ax647. After 1 h of incubation, hCOs were washed with PBS three times, fixed in 4% PFA in PBS at 4°C overnight, and immunostained. For ligand-directed chemical labeling for AMPARs and NMDARs (Nonaka et al., 2024; Wakayama et al., 2017), hCOs were reacted with 1 µM CAM2-Ax555 and 1 µM CNR1M-Ax647 in DMEM/F12 + Neurobasal medium for 1 h at 37°C under a humidified atmosphere of 5% CO₂. As a control for chemical labeling, hCOs were reacted with 1 µM NLC-Ax647, a control compound lacking a ligand moiety.

### Tissue clearing

hCOs were subjected to the CUBIC method (Susaki et al., 2015), followed by immunohistochemistry. Briefly, hCOs were fixed with 4% PFA in PBS at 4°C overnight. The next day, they were washed with PBS three times and incubated with CUBIC Reagent-1 diluted 2-fold with H₂O for 3–18 h before incubation with Reagent-1 overnight at 37°C. Organoids were washed with PBS three times for 90 min and stained with anti-GFP antibodies in PBS containing 0.1% Triton X-100 and 5% Blocking One for 2 days at 37°C. They were re-washed with PBS three times for 90 min and incubated with secondary antibodies for 2 days at 37°C. For refractive index matching, hCOs were incubated with CUBIC Reagent-2 diluted 2-fold for 3–18 h at room temperature, followed by Reagent-2 overnight at room temperature. Cleared hCOs were imaged using confocal microscopy (LSM800, Zeiss).

### Quantification and statistical analysis

Values are expressed as mean ± SEM unless otherwise stated. Statistical analyses were performed using R. Details of the statistical analyses are also given in the figure legends. Probability values < 5% were considered significant.

## Results

### hCOs derived from hiPSCs exhibited *in vivo*-like gene expression and cytoarchitecture

We used the SFEBq and spinning culture methods (Eiraku et al., 2008; Xiang et al., 2017) with some modifications to generate hCOs from hiPSCs (Figure 1A). Undifferentiated hiPSCs were dissociated into single cells and seeded in a low-adherent 96-well V-bottom plate at 10,000 cells per well. Y-27632 (50 µM) was added to inhibit cell death, and A-83-01 (500 nM) and LDN-193189 (100 nM) were added to promote a neuroectoderm fate. IWR-1 *endo* (2 µM) was administered to induce a forebrain fate. For days 10–18, bFGF (20 ng/mL) was added to promote progenitor cell proliferation (Figures 1B and S1A), resulting in larger organoids and higher expression of the telencephalic genes *FOXG1*, *EMX1*, and *EMX2* compared with a control in the absence of bFGF (Figures S1B and S1C).

**Figure 1.**
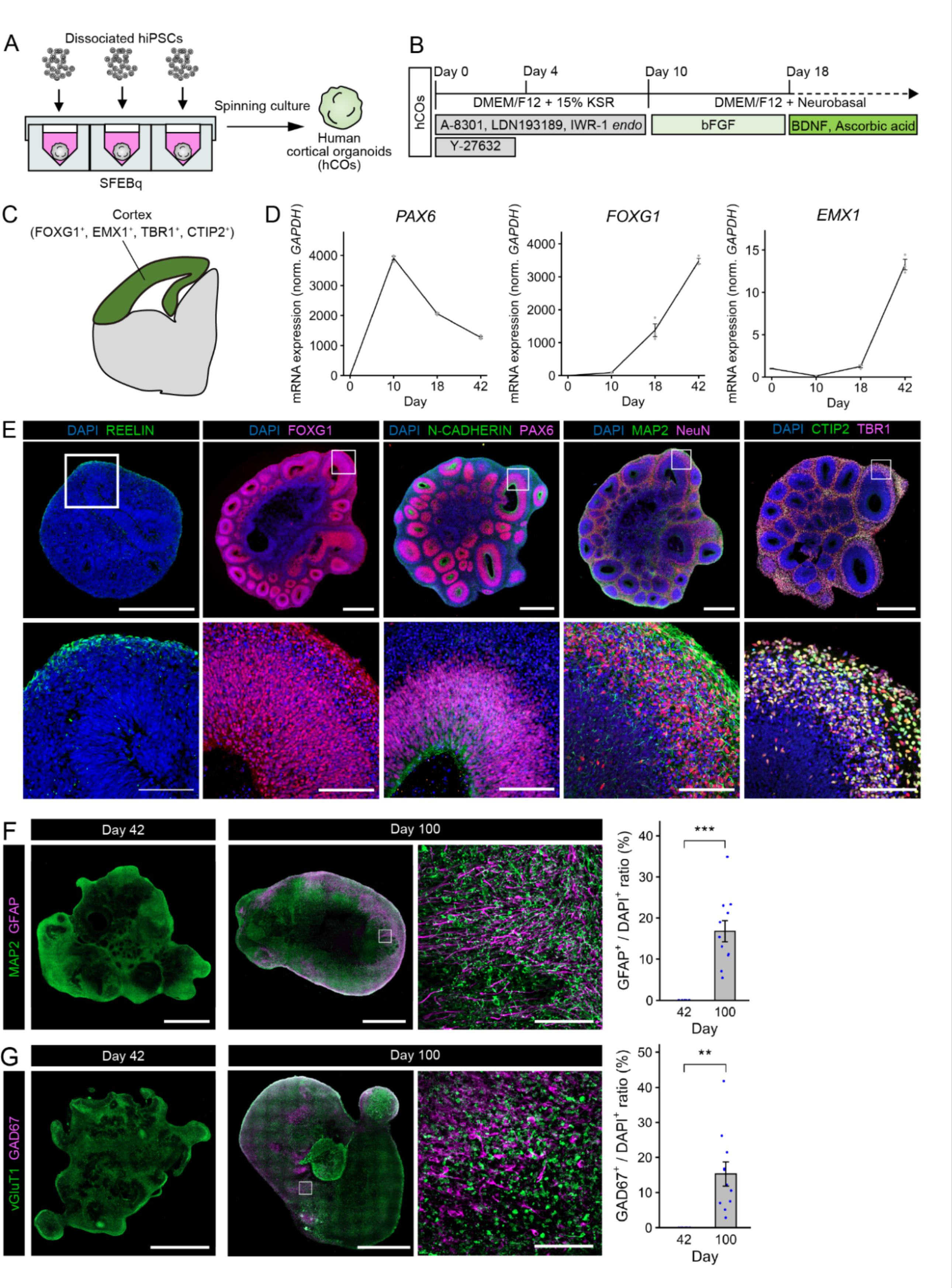
Human cortical organoids exhibited *in vivo*-like gene expression and cytoarchitecture. (A) Method for generating hCOs from hiPSCs. (B) Stepwise treatment for generating hCOs from hiPSCs. (C) Schematic representation of the expression patterns of cortical genes during development. (D) qPCR analysis of cortical markers in hCOs. Data are expressed as mean ± SEM; n = 3 organoids. (E) Immunostaining for REELIN in hCOs on day 18 and for FOXG1, N-CADHERIN, PAX6, MAP2, NeuN, TBR1, and CTIP2 in hCOs on day 42. Scale bars: 500 µm (top) and 100 µm (bottom). (F) Immunostaining for the neuronal marker MAP2 and the astrocyte marker GFAP and quantification of GFAP expression in hCOs on days 42 and 100. Data are expressed as mean ± SEM; n = 10 organoids (day 42) and n = 11 organoids (day 100). \*\*\**p* < 0.001, Welch’s *t*-test. (G) Immunostaining for the excitatory neuronal marker vGluT1 and the inhibitory neuronal marker GAD67 and quantification of GAD67 expression in hCOs on days 42 and 100. Data are expressed as mean ± SEM; n = 11 organoids (day 42) and n = 11 organoids (day 100). \*\**p* < 0.01, Welch’s *t*-test.

We first examined whether hCOs recapitulated cortical gene expression during brain development (Figure 1C). qPCR analysis in hCOs revealed increasing expression of the neural progenitor marker *PAX6* at an early stage (day 10), whereas the expression of *FOXG1* and *EMX1* increased over time (Figure 1D). To investigate hCO cytoarchitecture, we sectioned and analyzed them using immunohistochemistry. On day 18, FOXG1^+^ telencephalic cells and PAX6^+^ neural progenitors were observed in hCOs (Figure S1D). hCOs showed PAX6^+^ cells radially located outside the N-CADHERIN^+^ apical membrane, indicating neural tube-like structures or neural rosettes (Figure S1D). The immature neuronal marker TUJ1 was expressed in the outer layer of SOX2^+^ progenitors, indicating the adoption of an inside-out neurogenetic pattern in hCOs (Figure S1D). Additionally, REELIN^+^ Cajal– Retzius cells appeared in the superficial layer of hCOs (Figure 1E). On day 42, although FOXG1 was uniformly expressed, PAX6 expression became more restricted to neural rosettes compared with day 18 (Figure 1E). Furthermore, bFGF treatment promoted the formation of larger neural rosettes (Figure S1E). The mature neuronal markers MAP2 and NeuN were expressed outside neural rosettes, whereas the cortical excitatory neuron markers TBR1 and CTIP2 were detected in the outer neuronal layer (Figure 1E). These results show that hCOs mimicked the polarized cortical cytoarchitecture, including the ventricular zone and the cortical plate.

With cortical maturation, neurogenesis transitions to gliogenesis (Di Bella et al., 2024; Shen et al., 2006). Furthermore, unlike in the mouse brain, where all inhibitory neurons develop in the ganglionic eminence and migrate to the cerebral cortex, the inhibitory neurons of the human cortex are generated from both the ganglionic eminence and the cerebral cortex (Delgado et al., 2022). Therefore, we investigated whether glial cells and inhibitory neurons develop in hCOs over the course of culture maturation. Immunostaining showed that GFAP^+^ glial cells appeared in hCOs on day 100 but were absent on day 42 (Figure 1F). Additionally, GAD67^+^ inhibitory neurons were observed in hCOs on day 100 but not on day 42 (Figure 1G). Notably, expression of NKX2.1 and LHX6, inhibitory neuron markers derived from the medial ganglionic eminence, was not observed in hCOs on day 100 (Figures S2A and S2B). These results illustrate the transition from neurogenesis to gliogenesis and the generation of inhibitory neurons in hCOs.

### Neural activity in hCOs matured during development

The establishment of cortical circuits requires neural activity (Katz and Shatz, 1996; Luhmann et al., 2016). Therefore, we first investigated whether neurons in hCOs exhibit spontaneous neural activity. We infected the day 40 hCOs with AAVDJ-CaMKIIa-jGCaMP7f to label the excitatory neurons in hCOs with jGCaMP7f, a genetically encoded Ca^2+^ indicator (Dana et al., 2019). We then performed two-photon Ca^2+^ imaging 10 days after AAV infection (Figure 2A and Figure S3A). jGCaMP7f-labeled neurons in hCOs exhibited changes in jGCaMP7f fluorescence (Figure 2B). Action potentials are initiated by the opening of voltage-gated sodium channels in neurons. To determine whether the changes in jGCaMP7f fluorescence in hCOs were caused by action potentials, we examined the effect of TTX, a voltage-gated sodium channel blocker, on jGCaMP7f signals. The jGCaMP7f fluorescence changes in hCOs were blocked by TTX (1 μM), indicating the generation of spontaneous action potentials in the neurons of hCOs (Figures S3B and S3C). Next, we investigated changes in neural activity in hCOs over organoid culture maturation. We first isolated active cells among jGCaMP7f-labeled cells in the early-stage (day 50) and late-stage (day 80) hCOs and determined the frequency and amplitude of Ca^2+^ events for each active cell (Figure S4A). There was no significant difference in the active cell rate between days 50 and 80 (Figures 2C and 2D). Notably, hCOs on day 80 had Ca^2+^ events with higher frequency and larger amplitude than on day 50 (Figure 2D; Video S1), suggesting maturation of cortical neurons with spontaneous activity in late-stage hCOs.

**Figure 2.**
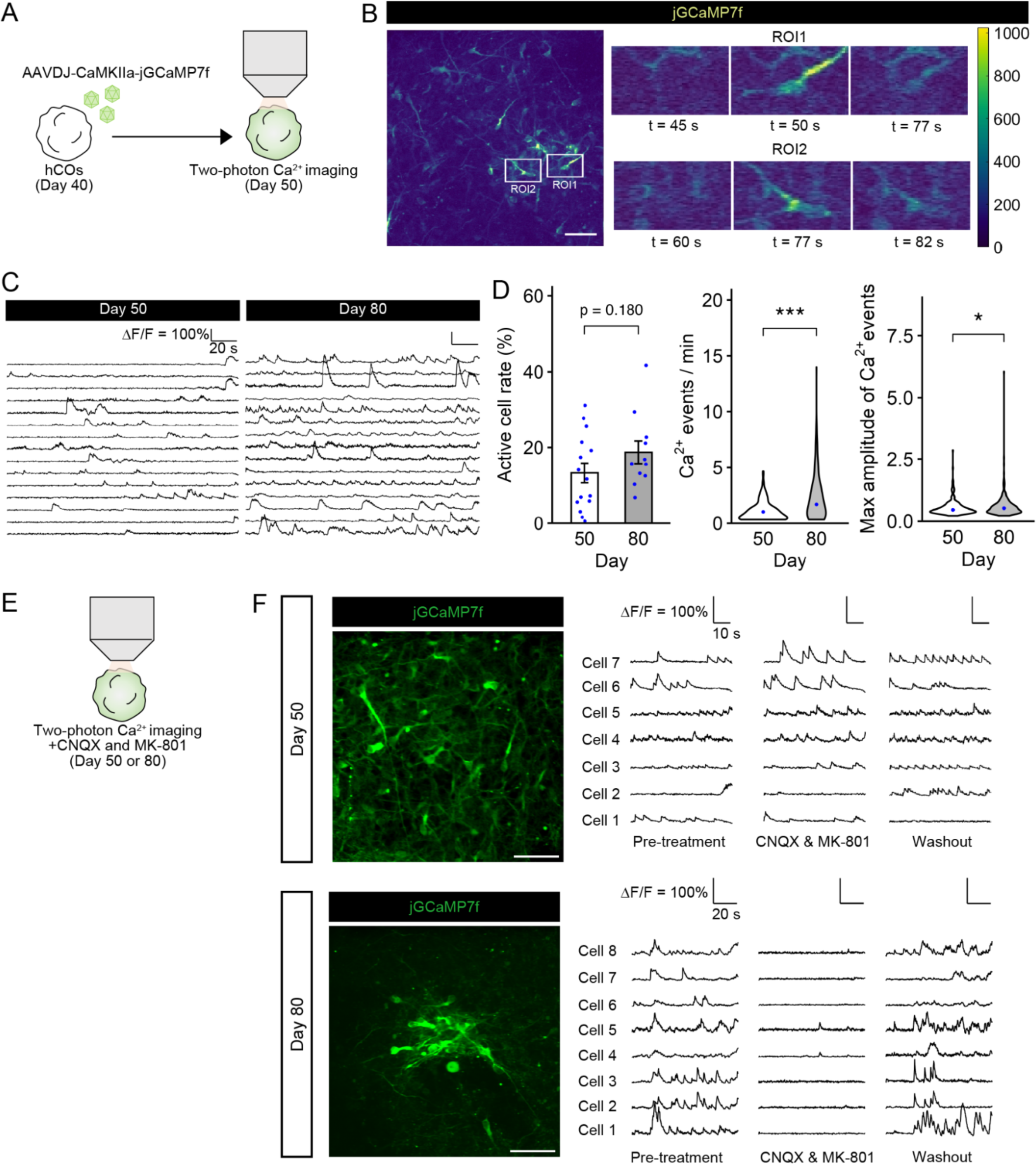
Neurons in human cortical organoids changed neural activity during development. (A) Schematic representation of viral labeling and two-photon Ca^2+^ imaging in hCOs. (B) Representative images showing jGCaMP7f fluorescent signal changes in individual neurons from hCOs on day 50. Scale bar: 50 µm. (C) Time-series traces of jGCaMP7f fluorescent signal changes in hCOs on days 50 and 80. (D) Developmental changes in neural activity in hCOs. Quantification of active cell rate: n = 15 organoids (day 50), n = 11 organoids (day 80), and Ca^2+^ events/min; n = 203 neurons (day 50), n = 255 neurons (day 80), and maximum amplitude of Ca^2+^ events; n = 203 neurons (day 50), n = 255 neurons (day 80) in hCOs. Data are expressed as mean ± SEM (active cell rate) and median (Ca^2+^ events/minute, amplitude of calcium events). Active cell rate: Welch’s *t*-test. Ca^2+^ events/minute, amplitude of calcium events: \**p* < 0.05, \*\*\**p* < 0.001, Mann–Whitney U test. (E) Two-photon Ca^2+^ imaging of hCOs in the absence or presence of CNQX (50 µM) and MK-801 (50 µM). (F) Effect of NMDAR and AMPAR antagonists on spontaneous neural activity in hCOs on days 50 (top) and 80 (bottom). Representative image of the imaging site and time-series traces of jGCaMP7f signal changes. Left: before application of CNQX and MK-801; middle: treatment of CNQX and MK-801; right: after washout of CNQX and MK-801. Scale bar: 50 µm.

To determine whether the observed neural activity in hCOs is mediated by excitatory synaptic transmission, we examined the effects of glutamate receptor antagonists on hCO neural activities during two-photon Ca^2+^ imaging (Figure 3E). Neural activities in hCOs on day 80 were blocked by CNQX (50 µM), an antagonist of AMPA and kainate receptors, and MK-801 (50 µM), an antagonist of NMDA receptors, whereas neural activities in hCOs on day 50 were not blocked (Figure 3F; Video S2). These results indicate that neural activities in late-stage hCOs, but not in early-stage hCOs, are mediated by glutamate transmission.

**Figure 3.**
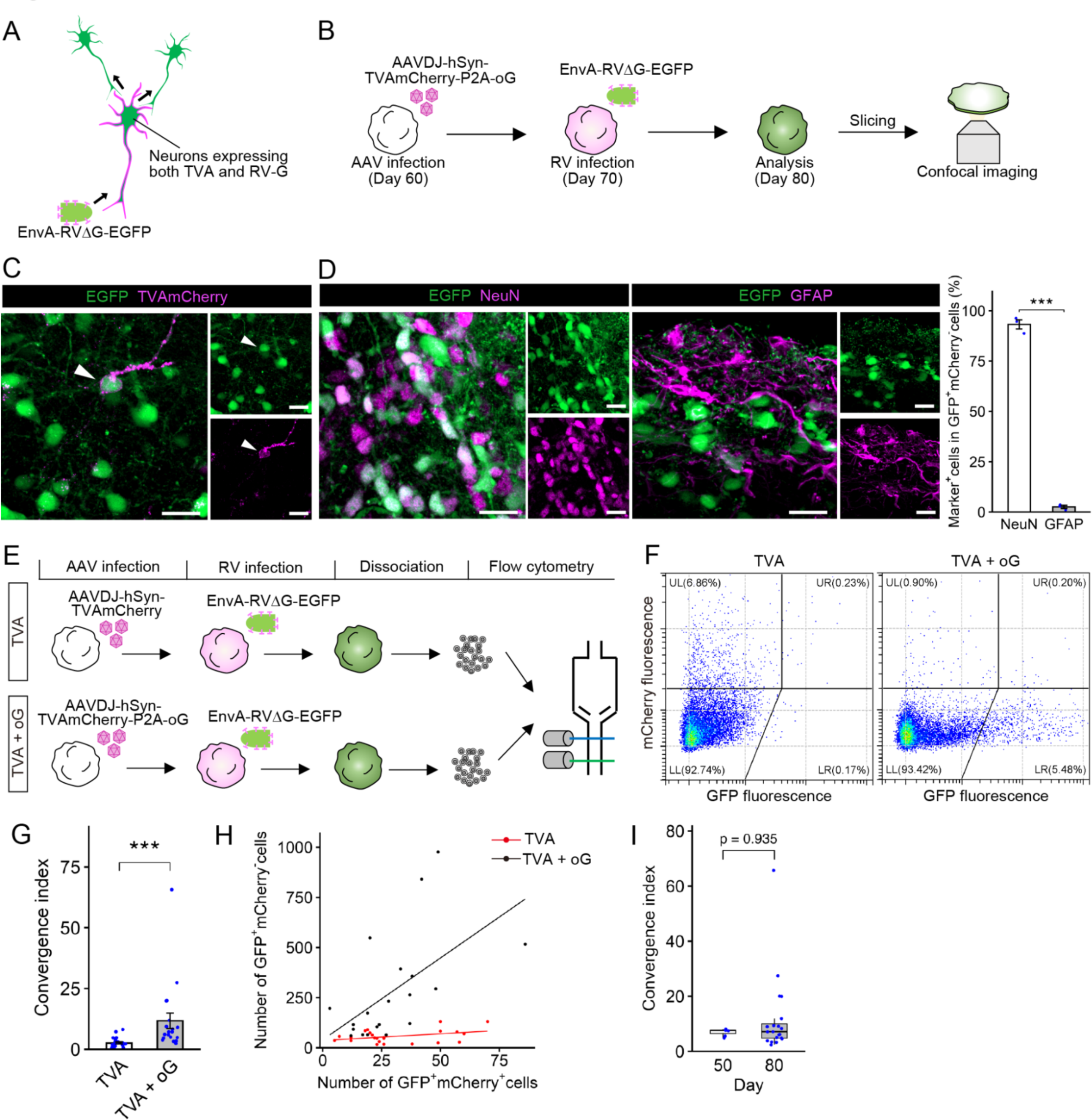
Neurons from human cortical organoids formed synaptic connections. (A) Monosynaptically restricted circuit tracing after targeted infection and trans-complementation of G-deleted rabies virus. (B) Experimental scheme for detection of synaptic connections in hCOs. (C) Immunostaining for mCherry and GFP in hCOs. Scale bar: 20 µm. White arrowhead, EGFP and mCherry double-positive neuron. (D) Representative images and quantification of the percentage of neurons (NeuN^+^) and glial cells (GFAP^+^) in retrogradely labeled cells (GFP^+^mCherry^−^). Scale bar: 20 µm. Data are expressed as mean ± SEM: n = 3 organoids. \*\*\**p* < 0.001, Welch’s *t*-test. (E) Experimental scheme for quantification of synaptic connections in hCOs. (F) Representative flow cytometry images of dissociated hCOs under TVA or TVA+oG conditions. (G) Convergence indices for hCOs under TVA or TVA+oG conditions. The convergence index was defined as the number of RVΔG-labeled input neurons divided by that of starter cells. Data are expressed as mean ± SEM: n = 23 organoids (TVA) and n = 20 organoids (TVA+oG). \*\*\**p* < 0.001, Mann–Whitney U test. (H) Scatter plot of the number of GFP^+^mCherry^+^ postsynaptic neurons and that of GFP^+^mCherry^−^ presynaptic cells under TVA or TVA+oG conditions. The plot represents an organoid (Pearson’s correlation coefficient value and *p*-value: r = 0.38, *p* = 0.07 in TVA. r = 0.57, *p* = 0.008 in TVA+oG). (I) Box plot of the convergence index for hCOs on days 50 and 80. n = 7 organoids (day 50) and n = 20 organoids (day 80).

Previous studies have shown that neural activity in the developing cortex changes from an asynchronous pattern to synchronized pattern (Molnár et al., 2020; Wu et al., 2024). To evaluate the activity synchronization of cortical neurons in hCOs at the population level, we calculated the mean correlation coefficients between active cell pairs. The correlation between neural activities was low in hCOs on days 50 and 80, indicating that the neural activity on both days was mostly asynchronous with a low probability of synchronized activity (Figure S4B). We also performed PCA to determine the variability in neural activity patterns in hCOs. The first principal component explained approximately 40% of the total variance on both days 50 and 80, indicating equally diverse activity patterns in hCOs between days 50 and 80 (Figures S4C and S4D). Thus, over hCO development, neural activity in hCOs matured *per se*, but did not elicit synchronous spontaneous activity in the population.

### Neurons in hCOs formed synaptic connections

To evaluate synaptic formation in hCOs, we analyzed hCOs by immunohistochemistry for synaptic proteins. On day 72, the excitatory presynaptic and postsynaptic markers vGluT1 and HOMER1, respectively, colocalized on MAP2^+^ neurites of neurons in hCOs, suggesting formation of synaptic contacts between presynaptic excitatory neurons and their targets (Figure S5). To elucidate the synaptic connectivity within hCOs, we performed retrograde transsynaptic tracing using RVΔG (Osakada et al., 2011; Wickersham et al., 2007) (Figure 3A). In this approach, presynaptic neurons were transsynaptically infected with RVΔG from postsynaptic neurons expressing both TVA and optimized glycoprotein (oG) (Kim et al., 2016) following targeted infection with EnvA-pseudotyped RVΔG and trans-complementation with G. On day 60, we first infected hCOs with AAVDJ-hSyn-TVAmCherry-P2A-oG, which encodes the TVA receptor and RV-G protein required for targeted infection and transsynaptic spread of EnvA-pseudotyped RVΔG, respectively. Subsequently, hCOs were infected with EnvA-pseudotyped RVΔG encoding EGFP (EnvA-RVΔG-EGFP). Ten days after rabies viral infection, we sectioned hCOs and performed immunohistochemistry (Figure 3B). Both EGFP and mCherry double-positive cells and EGFP single-positive cells were observed in hCOs (Figure 3C). EGFP single-positive cells indicate RVΔG transsynaptic spread from EGFP and mCherry double-positive starter cells to their direct presynaptic inputs. Although most EGFP^+^ mCherry^−^ cells expressed the neuronal marker NeuN, a few populations expressed the glial marker GFAP (Figure 3D). These results demonstrate formation of synaptic connections between neurons in hCOs.

We quantitatively analyzed the synaptic connections in hCOs by conducting transsynaptic rabies tracing and flow cytometry and counting the number of EGFP^+^mCherry^+^ postsynaptic neurons and EGFP^+^mCherry^−^ presynaptic neurons (Figure 3E). In the control condition (TVA alone), hCOs were infected with AAVDJ-hSyn-TVAmCherry, followed by EnvA-RVΔG-EGFP. Despite the presence of EGFP^+^mCherry^+^ cells, few EGFP^+^ mCherry^−^ cells were observed in the control. In contrast, in the presence of oG (TVA + oG), the number of EGFP^+^ mCherry^−^ cells significantly increased, indicating transsynaptic viral spread from starter cells (Figure 3F). The convergence index, defined as the ratio of EGFP^+^mCherry^−^ presynaptic cells to EGFP^+^mCherry^+^ postsynaptic cells (Kim et al., 2016), was significantly higher in hCOs expressing both TVA and oG than in those expressing TVA alone (Figure 3G). Furthermore, in the presence of oG, the number of postsynaptic neurons increased along with the number of presynaptic neurons (Figure 3H). Notably, the convergence index did not differ between days 50 and 80 hCOs (Figure 3I). These results indicate that neurons in hCOs formed synaptic connections at similar levels on days 50 and 80.

### AMPAR expression on the cell membrane increased during developmental processes in hCOs

Synaptic transmission via the neurotransmitter glutamate in the mammalian cortex is primarily mediated by NMDARs and AMPARs (Chater and Goda, 2014). To investigate the developmental expression of these receptors in hCOs, we performed qPCR analysis for NMDAR and AMPAR subunit genes at early (days 45–50) and late (days 75–90) stages. The mRNA expression levels of the NMDAR subunits *GRIN1*, *GRIN2A*, and *GRIN2B* and the AMPAR subunits *GRIA1* and *GRIA2* at the early stage were comparable to those at the late stage (Figures S6A and S6B). To visualize and quantify the AMPAR protein distributed in the cell membrane, we used ligand-directed chemical labeling for hCOs (Wakayama et al., 2017). The CAM2-Ax647 reagent, which consists of the AMPAR ligand PFQX and the fluorescent dye Alexa647, enables AMPAR visualization on the cell membrane using Alexa647 through a chemical reaction mediated by ligand–receptor binding (Figures 4A–C). Alexa647-labeled puncta were observed along the MAP2^+^ membrane in hCOs treated with CAM2-Ax647, but none were detected in control hCOs treated with the no-ligand reagent NLC-Ax647. Moreover, these puncta significantly decreased in the presence of CNQX, a competitive AMPAR antagonist, thereby verifying specific AMPAR labeling by CAM2-Ax647 (Figures 4D and 4E). Using this method, we next compared AMPAR expression between days 50 and 80 hCOs and found a significant increase in the number of Alexa647-labeled puncta on day 80 (Figures 4F and 4G). Notably, these puncta were colocalized with the excitatory postsynaptic marker HOMER1 (Figure S7), suggesting that hCOs acquired AMPAR-localized synapses over maturation.

**Figure 4.**
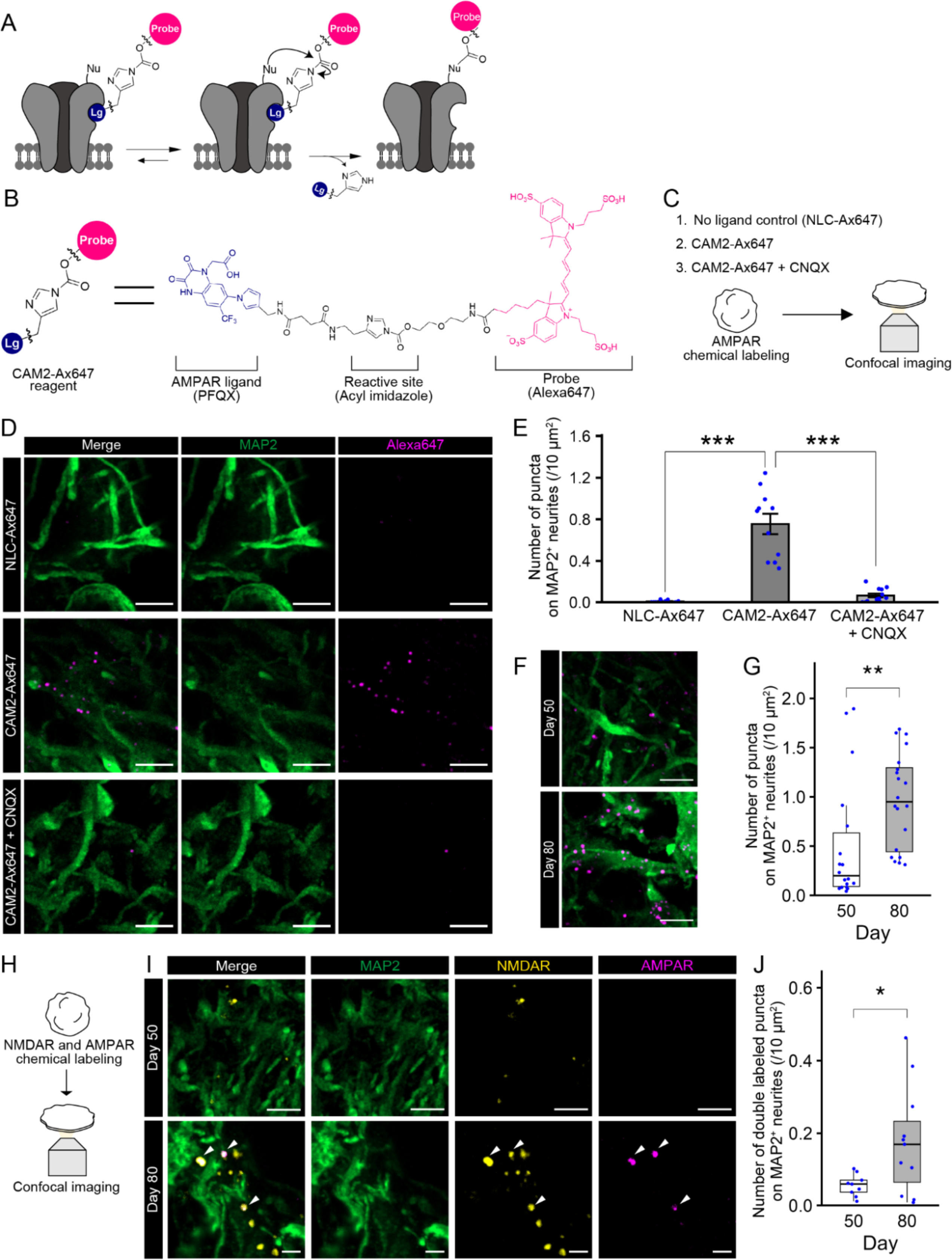
AMPA receptor expression on the cell membrane in human cortical organoids increased over development. (A) Ligand-directed chemical labeling for AMPA receptors. (B) Chemical structure of CAM2-Ax647. Note that the hydrophilic and cell-impermeable features of CAM2-Ax647 suppress nonspecific adsorption to hydrophobic materials in tissues and promote covalent labeling of only cell-surface receptors. (C) Chemical labeling and AMPAR analysis in hCOs. (D) Immunostaining for MAP2 on day 80 hCOs treated with NLC-Ax647, CAM2-647, or CAM2-647+CNQX. Scale bar: 5 µm. (E) Quantification of the number of Alexa647-labeled puncta on MAP2^+^ neurites for each condition. Data are expressed as mean ± SEM; n = 12 organoids (NLC-Ax647 and CAM2-Ax647+CNQX) and n = 11 organoids (CAM2-Ax647). \*\*\**p* < 0.001, Tukey’s test. (F) Immunostaining for MAP2 on days 50 and 80 hCOs treated with CAM2-Ax647. Scale bar: 5 µm. (G) Quantification of the number of Alexa647-labeled puncta on MAP2^+^ neurites in hCOs on days 50 and 80. n = 18 organoids (day 50) and n = 20 organoids (day 80). \*\**p* < 0.01, Mann–Whitney U test. (H) Chemical labeling for NMDAR and AMPAR in hCOs. (I) Immunostaining for MAP2 on days 50 and 80 hCOs treated with CNR1M-Ax647 and CAM2-Ax555. Scale bar: 2 µm. White arrowheads, colocalization of Alexa647 (NMDAR) and Alexa555 (AMPAR) on MAP2^+^ neurites. Scale bar: 5 µm (top) and 2 µm (bottom). (J) Quantification of the number of Alexa555 and Alexa647 double-labeled puncta on MAP2^+^ neurites in hCOs on days 50 and 80. n = 9 organoids (day 50) and n = 11 organoids (day 80). \**p* < 0.05, Welch’s *t*-test.

Synapses are constantly generated in the developing brain. However, newly generated glutamatergic synapses lack functional AMPAR-mediated transmission. During development, silent synapses expressing only NMDARs are converted to functional synapses containing both NMDARs and AMPARs (Hanse et al., 2013; Kanold et al., 2019). To investigate whether synapses in late-stage hCOs contain NMDAR and AMPAR, we conducted chemical labeling for NMDARs and AMPARs using CNR1M-Ax647 (NMDAR) and CAM2-Ax555 (AMPAR) on days 50 and 80 (Figure 4H and Figures S8A–D). We observed a significant increase of colocalization of Alexa647-labeled puncta (NMDAR) and Alexa555-labeled puncta (AMPAR) on day 80, suggesting that synapses in late-stage hCOs contain both NMDARs and AMPARs (Figures 4I and 4J). Based on our results, we conclude that hCOs recapitulate the conversion of silent synapses to functional synapses.

## Discussion

In the present study, we examined developmental changes in neural activity and synaptic connectivity in hCOs. Two-photon Ca^2^⁺ imaging showed more mature neural activity by day 80 compared to day 50. RVΔG circuit tracing revealed similar synaptic connectivity on both days, whereas chemical labeling indicated a significant increase in AMPAR expression on the cell membrane as well as colocalization with NMDARs at the late stage. Pharmacological perturbations confirmed AMPAR-mediated functional synaptic transmission in late-stage hCOs. Thus, we conclude that hiPSC-derived neural organoids progressively organize AMPAR-mediated excitatory synaptic transmission in cortical circuits along a developmental trajectory.

In the developing cerebral cortex, polarized compartments, such as the ventricular zone and cortical plate, give rise to diverse cell populations and a six-layer cortical structure, with gliogenesis occurring after the peak of neurogenesis (Greig et al., 2013; Kelley and Pașca, 2022). We found that hCOs formed distinct ventricular zone-like and cortical plate-like compartments. In addition, hCOs not only recapitulated the transition from neurogenesis to gliogenesis but also reflected the primate-specific production of inhibitory neurons from cortical progenitors (Delgado et al., 2022). Thus, hCOs closely mimic the development of cellular identity and cytoarchitecture in the human cerebral cortex.

During early brain development, synaptic connections are initially silent, becoming active through the recruitment of AMPARs on the postsynaptic membrane (Hanse et al., 2013; Kanold et al., 2019). In this study, RVΔG tracing revealed similar numbers of synaptic connections in hCOs between days 50 and 80. However, chemical labeling showed a significant increase in AMPAR expression and colocalization with NMDARs by day 80. Additionally, two-photon Ca^2^⁺ imaging with AMPAR/NMDAR antagonists indicated functional maturation of AMPAR-dependent synaptic transmission on day 80. These findings suggest that although most synapses in hCOs were silent on day 50, they became active with increased AMPAR expression on the postsynaptic membrane by day 80. Thus, hCOs exhibit a developmental trajectory of physiological maturation characterized by the conversion of silent synapses into functional synapses and establishment of excitatory synaptic transmission, closely recapitulating *in vivo* cortical circuit development.

Synaptogenesis is a continuous process in the developing brain, and disruptions in this process are linked to various neurodevelopmental disorders (Betancur et al., 2009). Thus, evaluating dynamic changes in human synaptogenesis is essential for understanding these mechanisms. In this study, we developed a system to detect and quantify synaptic connections in hCOs. Traditional methods rely on the colocalization of synaptic proteins via immunohistochemistry, which can lead to false-positive synaptic connections due to incidental colocalization. Our approach using RVΔG, AAV, and cell sorting technology addresses this issue, enabling precise quantification of synapses in hCOs. This system facilitates detailed investigations of synaptic changes during development and pathology. A previous study indicates that the efficiency of RVΔG circuit tracing is unaffected by cortical cell type or synapse location (Patiño et al., 2023). In addition, our results suggest that rabies virus can spread across both silent and functional synapses and that RVΔG circuit tracing detects synaptic structures and labels presynaptic cells, regardless of the functional state of the synapse.

In the developing cortex, spontaneous neural activity transitions from asynchronous to synchronous patterns, which are critical for shaping cytoarchitecture, circuit organization, and area patterning (Molnár et al., 2020; Wu et al., 2024). An intriguing question is whether hCOs can generate synchronous neural activity. Although hCOs displayed spontaneous activity and functional maturation of synaptic transmission through developmental processes, synchronous activity was rarely observed even at later stages. In rodents, wave-like synchronous activity originates in the thalamus, which acts as a hub connecting peripheral tissues to the cortex, propagating waves that trigger synchronous cortical activity (Moreno-Juan et al., 2017). The absence of thalamic inputs in isolated hCOs may account for the limited synchronous activity observed in our system. A future challenge is to establish assembloids through the fusion of cortical organoids and thalamic organoids to recapitulate synchronous activity patterns in the human cerebral cortex.

In conclusion, this study demonstrated the developmental changes in AMPAR-mediated synaptic transmission in human cortical circuits using neural organoid technology. hCOs derived from hiPSCs replicate *in vivo* developmental programs essential for physiological maturation, offering a valuable model for studying human corticogenesis and related neurodevelopmental disorders.

## Abbreviations

hCO: human cortical organoid
AMPAR: α-amino-3-hydroxy-5-methyl-4-isoxazolepropionic acid receptor
NMDAR: N-methyl-D-aspartate receptor
hiPSC: human induced pluripotent stem cell
RVΔG: G-deleted rabies virus vector
AAV: adeno-associated vector

## Acknowledgments

We thank members of the Osakada Laboratory for their valuable discussions; Mr. Hanada, Mr. Nishimura, Mr. Kano, Mr. Kato, and Mr. Kobayashi (Research Equipment Development Group, Technical Center, Nagoya University) for manufacturing instruments.

## Author contributions

M.N. cultured hiPSCs, generated organoids, performed virus production, cell sorting, rabies tracing experiments, and imaging experiments, collected and analyzed all data, and wrote the manuscript. T.K. contributed to hiPSCs culture, organoids generation, cell sorting, and imaging experiments. S.A. cultured hiPSCs, generated organoids, and contributed to virus production. A.Y.S. and R.F.T. contributed to imaging data analysis. H.N. and I.H. synthesized AMPAR and NMDAR labeling reagents. F.O. wrote the manuscript and supervised the project.

---

**Figure S1.**
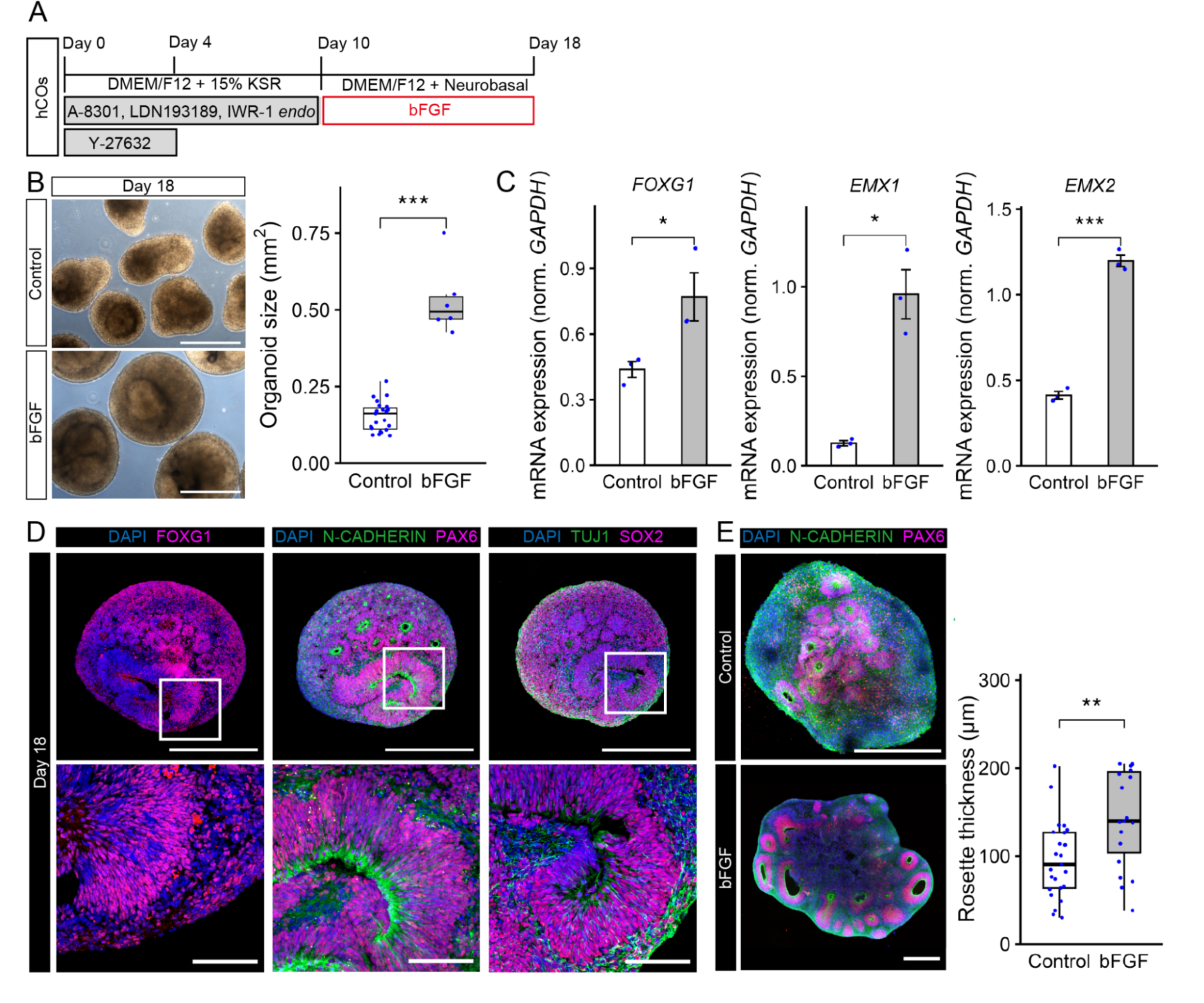
Effect of bFGF on hCO generation, related to Figure 1. (A) Protocol for generating hCOs from hiPSCs in the absence/presence of bFGF. (B) Bright-field images of hCOs (left) and quantification of hCO size in the absence or presence of bFGF on day 18 (right). Scale bar: 500 µm. n = 23 organoids (Control) and n = 6 organoids (bFGF). \*\*\**p* < 0.001, Welch’s *t*-test. (C) qPCR analysis for cortical markers in hCOs with/without bFGF. Data are expressed as mean ± SEM. \**p* < 0.05, \*\*\**p* < 0.001, Welch’s *t*-test. (D) Immunostaining for FOXG1, N-CADHERIN, PAX6, TUJ1, and SOX2 in hCOs with bFGF on day 18. Scale bars: 500 µm (top) and 100 µm (bottom). (E) Immunostaining and quantification of neural rosette thickness in hCOs in the absence/presence of bFGF on day 42. Scale bar: 500 µm. n = 25 rosettes (Control) and n = 19 rosettes (bFGF), \*\**p* < 0.01, Mann–Whitney U test.

**Figure S2.**
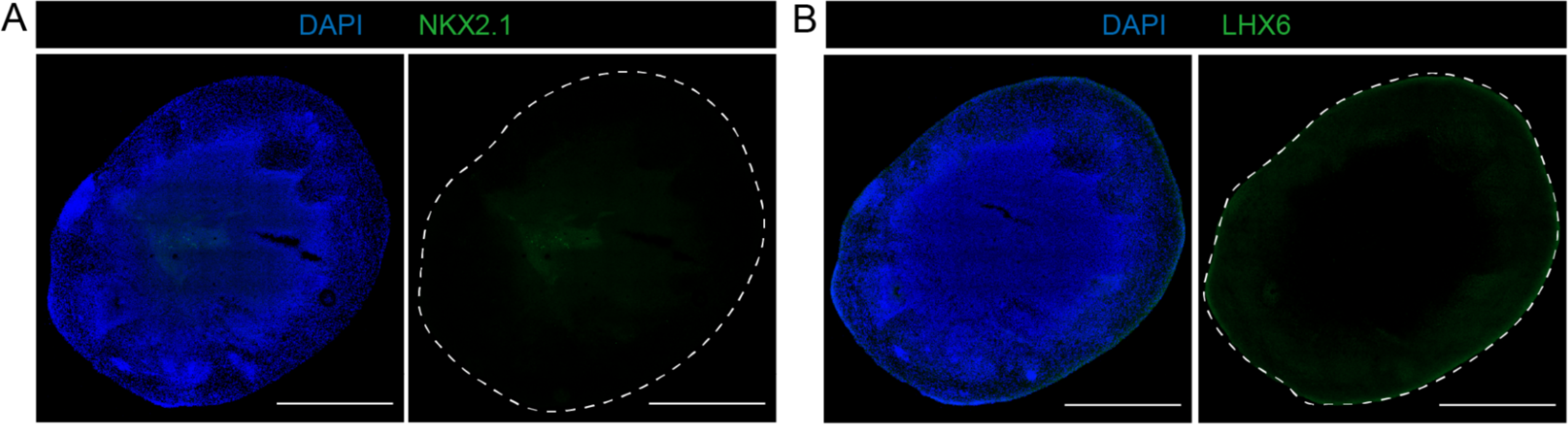
Missing inhibitory neuron lineage derived from the ganglionic eminence in hCOs, related to Figure 1. (A) Immunostaining for NKX2.1, a marker for inhibitory neurons derived from the ganglionic eminence, in hCOs on day 100. Scale bar: 1000 µm. (B) Immunostaining for LHX6, a marker for inhibitory neurons derived from the ganglionic eminence, in hCOs on day 100. Scale bar: 1000 µm.

**Figure S3.**
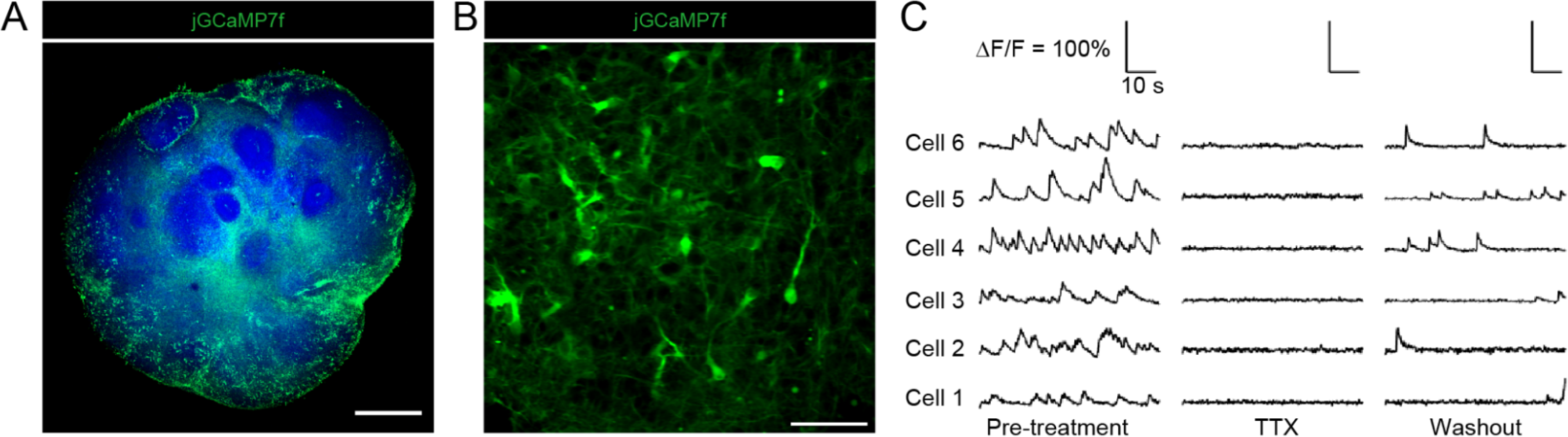
TTX application in hCOs, related to Figure 2. (A) Three-dimensional immunostaining for jGCaMP7f in hCOs on day 50. Scale bar: 500 µm. (B) Representative image of the imaging site on day 50 hCOs. Scale bar: 50 µm. (C) Effect of TTX, a voltage-gated sodium channel antagonist, on spontaneous neural activity in hCOs on day 50. Time-series traces of jGCaMP7f signal changes. Left: before application of TTX; middle: treatment of TTX; right: after washout of TTX. Scale bar: 50 µm.

**Figure S4.**
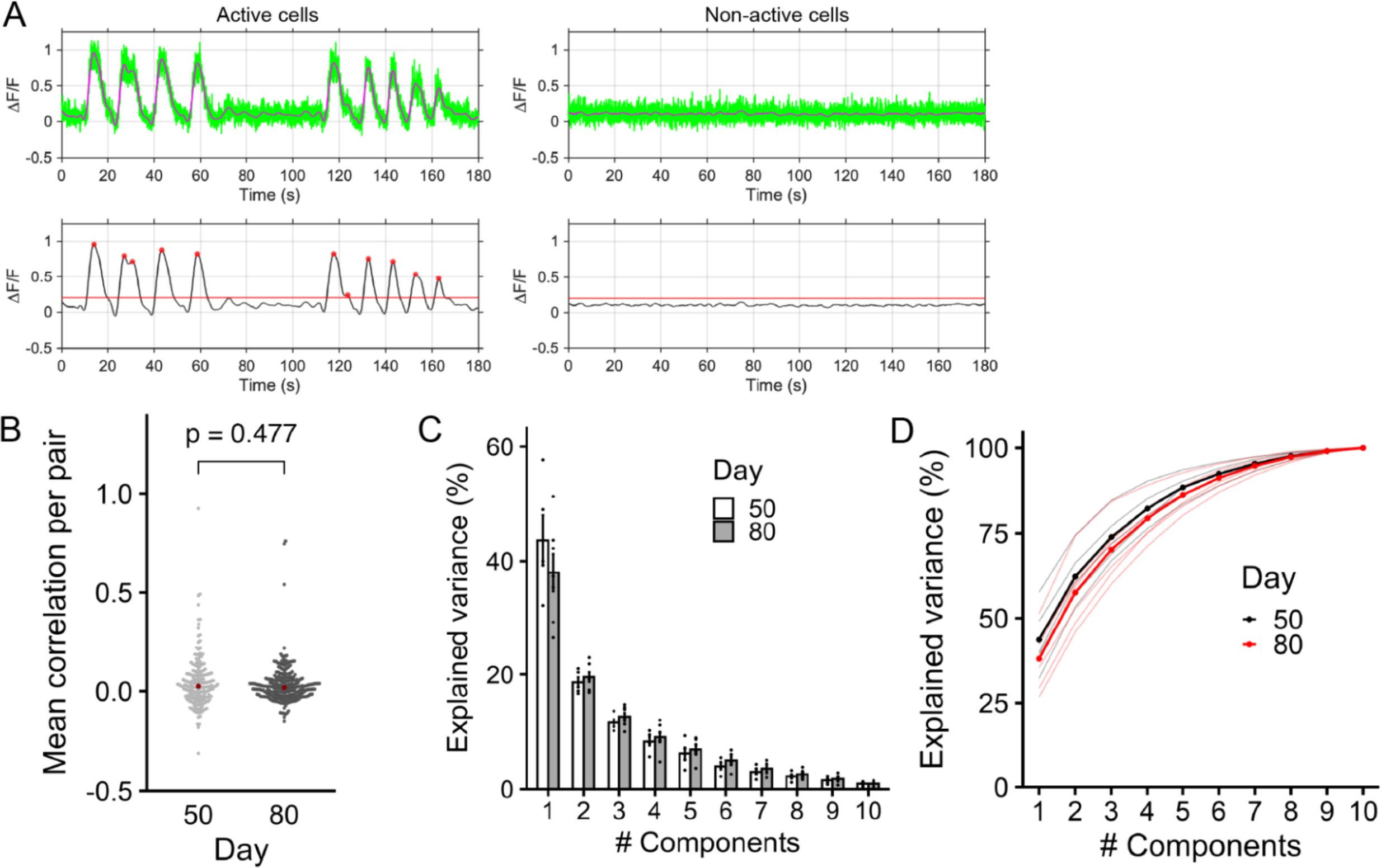
Analysis of Ca^2+^ imaging data from hCOs, related to Figure 2. (A) Representative time-series traces of jGCaMP7f fluorescent signal changes in active and non-active cells. Top: Green lines, raw signals; magenta lines, filtered signals. Bottom: Red lines, threshold that determines whether the fluorescent signal change is a calcium event; red dots, detected calcium events. (B) Mean correlation coefficients of activity between all pairs of simultaneously recorded neurons from hCOs on days 50 and 80. Gray points, correlation of each cell pair; red points, median. Mann–Whitney U test. (C) Percentages of explained variance for each principal component value on days 50 and 80 hCOs on PCA; n = 5 imaging sites (day 50), n = 7 imaging sites (day 80). (D) Cumulative explained variance plot for days 50 and 80 hCOs on PCA. Two solid bold lines, mean; black or red thin lines; n = 5 imaging sites (day 50), n = 7 imaging sites (day 80).

**Figure S5.**
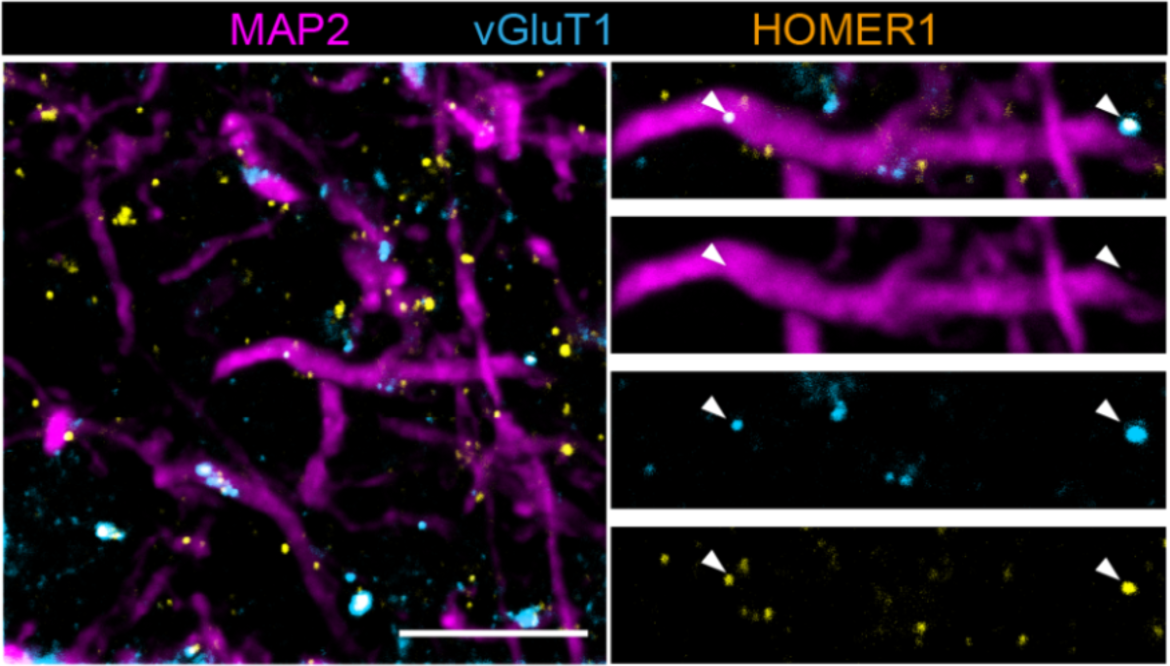
Formation of excitatory synapses in hCOs, related to Figure 3. Immunostaining for MAP2 (neurite marker), vGluT1 (presynaptic marker), and HOMER1 (postsynaptic marker). Scale bar: 10 µm. White arrowheads, vGluT1 and HOMER1 colocalization on MAP2^+^ neurites.

**Figure S6.**
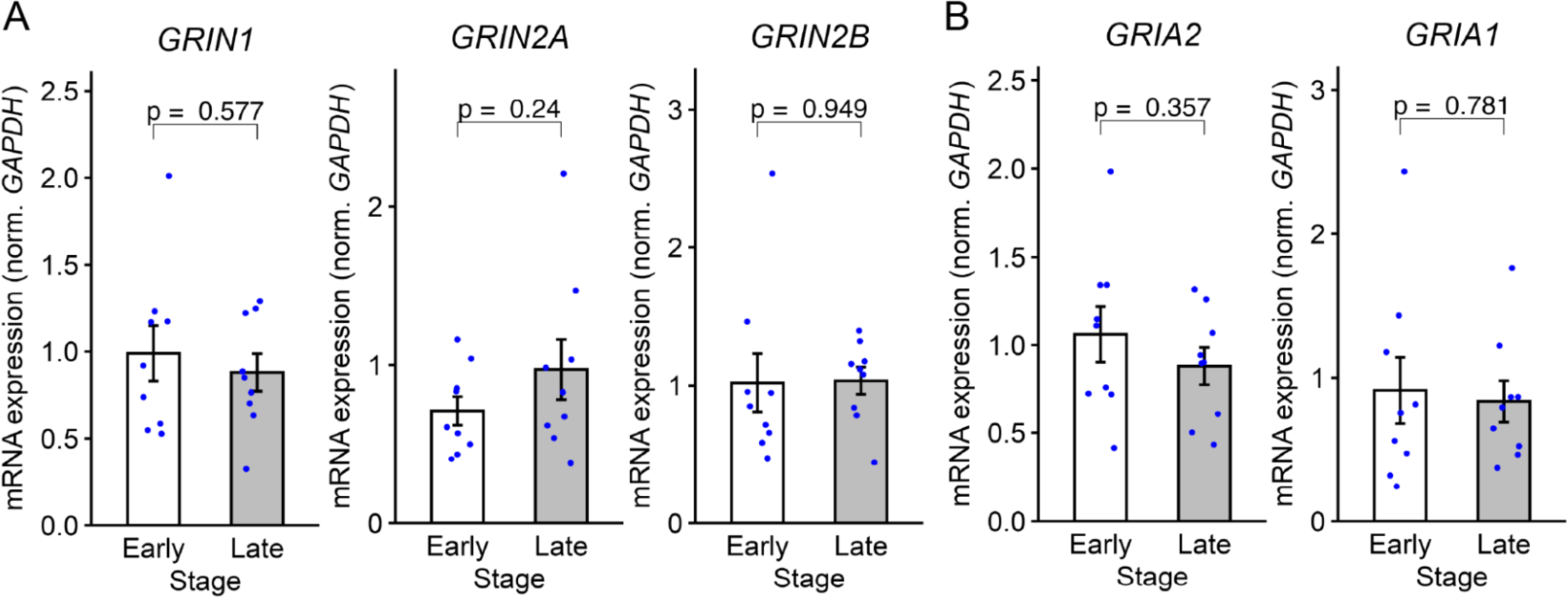
qPCR analysis of glutamate receptor expression during hCO development, related to Figure 4. (A) qPCR for *GRIN1*, *GRIN2A*, and *GRIN2B*, NMDAR subunits. Early and late stages represent hCOs on days 45–50 and 75-90, respectively. Data are expressed as mean ± SEM. n = 9 organoids. Welch’s *t*-test. (B) qPCR for *GRIA1* and *GRIA2*, AMPAR subunits. Early and late stages represent hCOs on days 45–50 and 75-90, respectively. Data are expressed as mean ± SEM. n = 9 organoids. Welch’s *t*-test.

**Figure S7.**
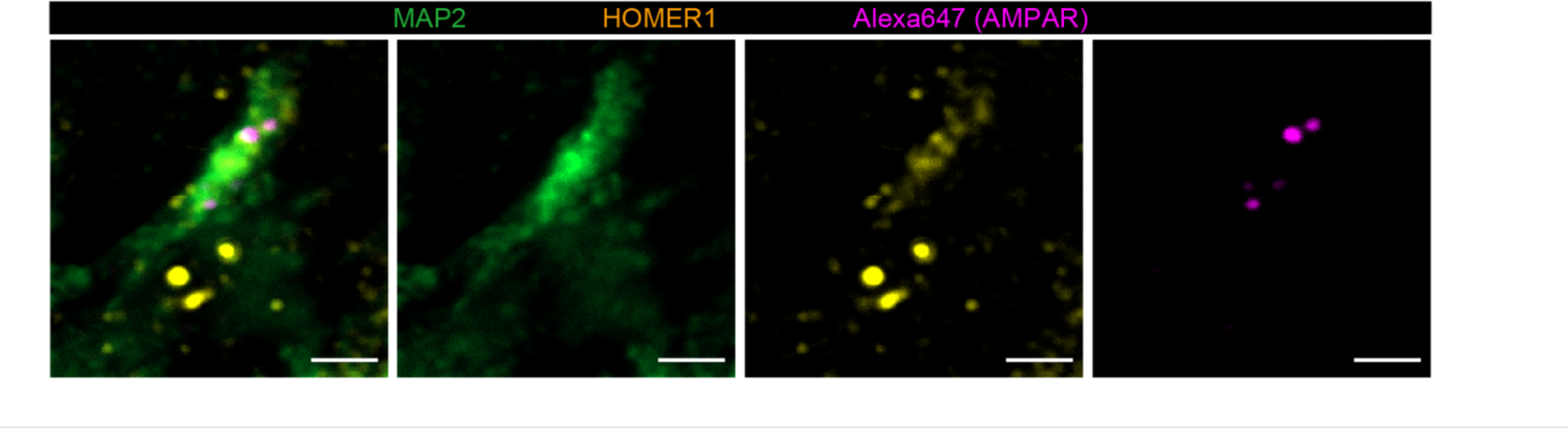
Distribution of AMPRARs to postsynapse in hCOs, related to Figure 4. Immunostaining for MAP2 (neurite marker) and HOMER1 (postsynaptic marker) on day 80 hCOs treated with CAM2-Ax647. Scale bar: 2 µm.

**Figure S8.**
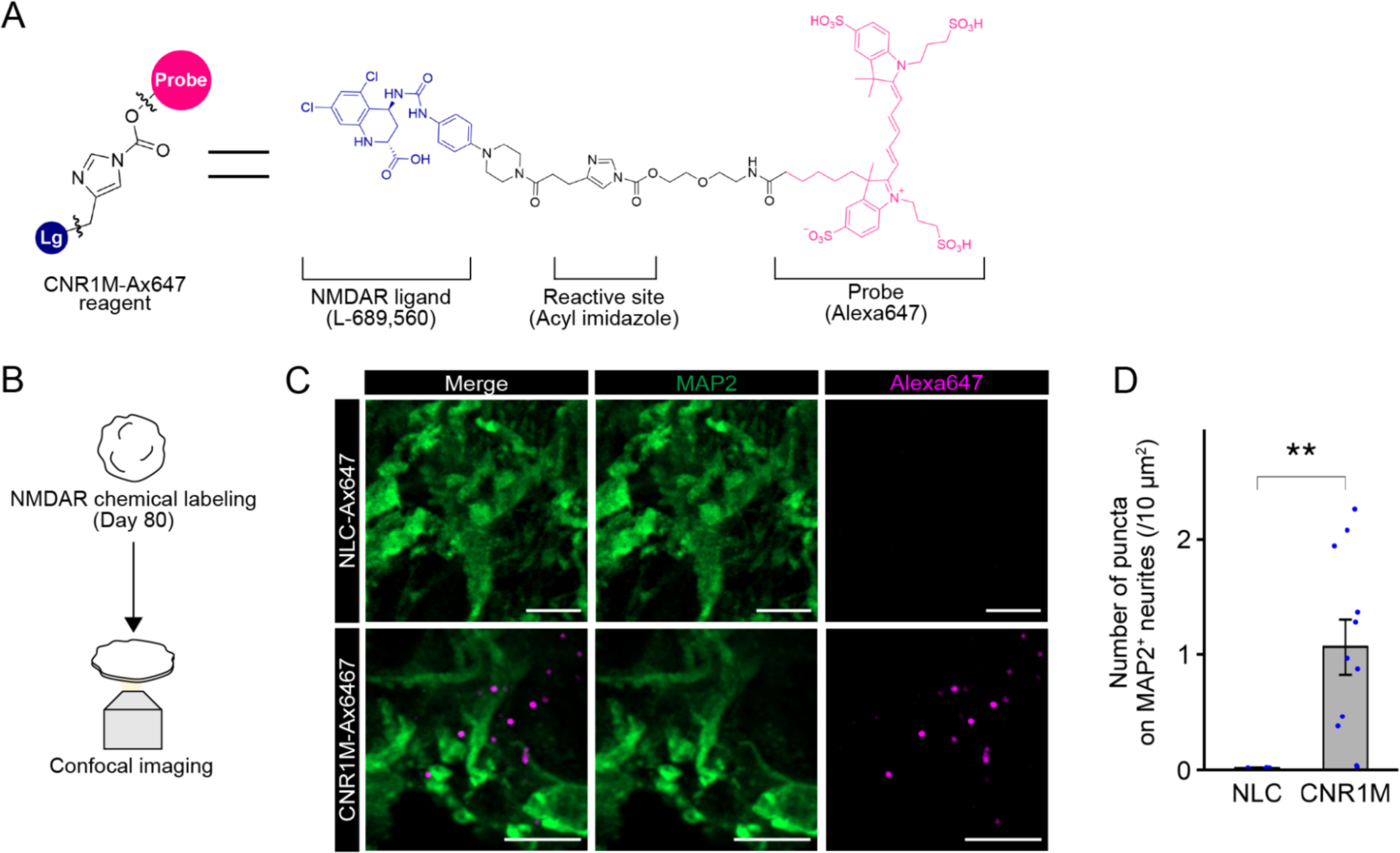
Chemical labeling for NMDARs with CNR1M-Ax647, related to Figure 4. (A) Chemical structure of CNR1M-Ax647. (B) Schematic of chemical labeling of NMDAR in hCOs. (C) Immunostaining for MAP2 on day 80 hCOs treated with NLC-Ax647 or CNR1M-Ax647. Scale bar: 5 µm. (D) Quantification of the number of Alexa647-labeled puncta on MAP2^+^ neurites in hCOs treated with NLC-Ax647 or CNR1M-Ax647. Data are expressed as mean ± SEM; n = 4 organoids (NLC-Ax647) and n = 11 organoids (CNR1M-Ax647). \*\**p* < 0.01, Welch’s *t*-test.

## Supplementary movie 1

Two-photon Ca^2+^ imaging of spontaneous neural activity in early-stage and late-stage hCOs. Two-photon imaging was performed on jGCaMP7f-labeled hCOs on days 50 and 80.

## Supplementary movie 2

Two-photon Ca^2+^ imaging of glutamate receptor-mediated neural activity in late-stage hCOs.

Two-photon imaging on jGCaMP7f-labeled hCOs on day 80 before (first) and during (second) CNQX (50 µM) and MK-801 (50 µM) application and after washout (third).

## Supplementary Information

### Primers for qPCR analysis

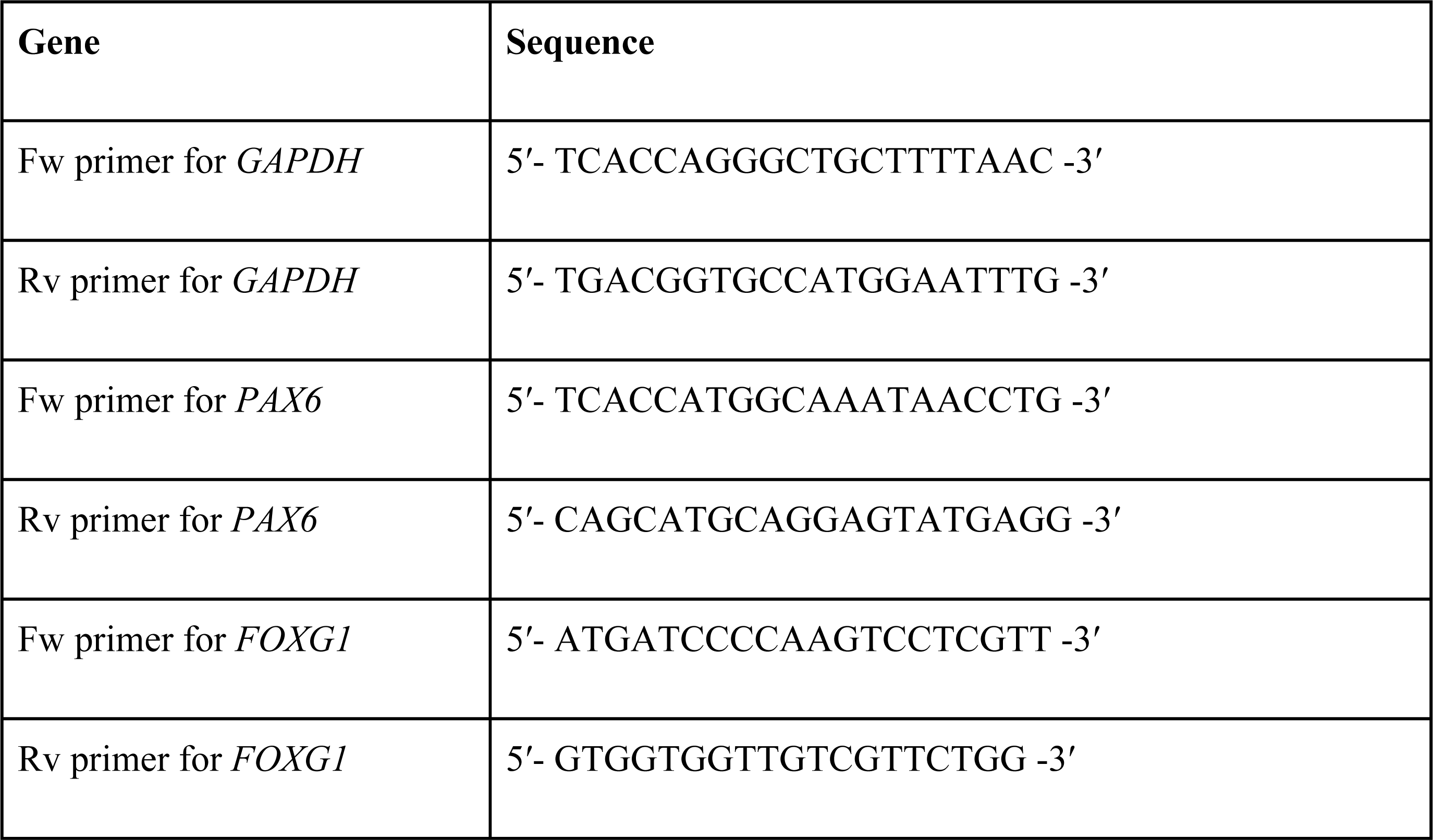
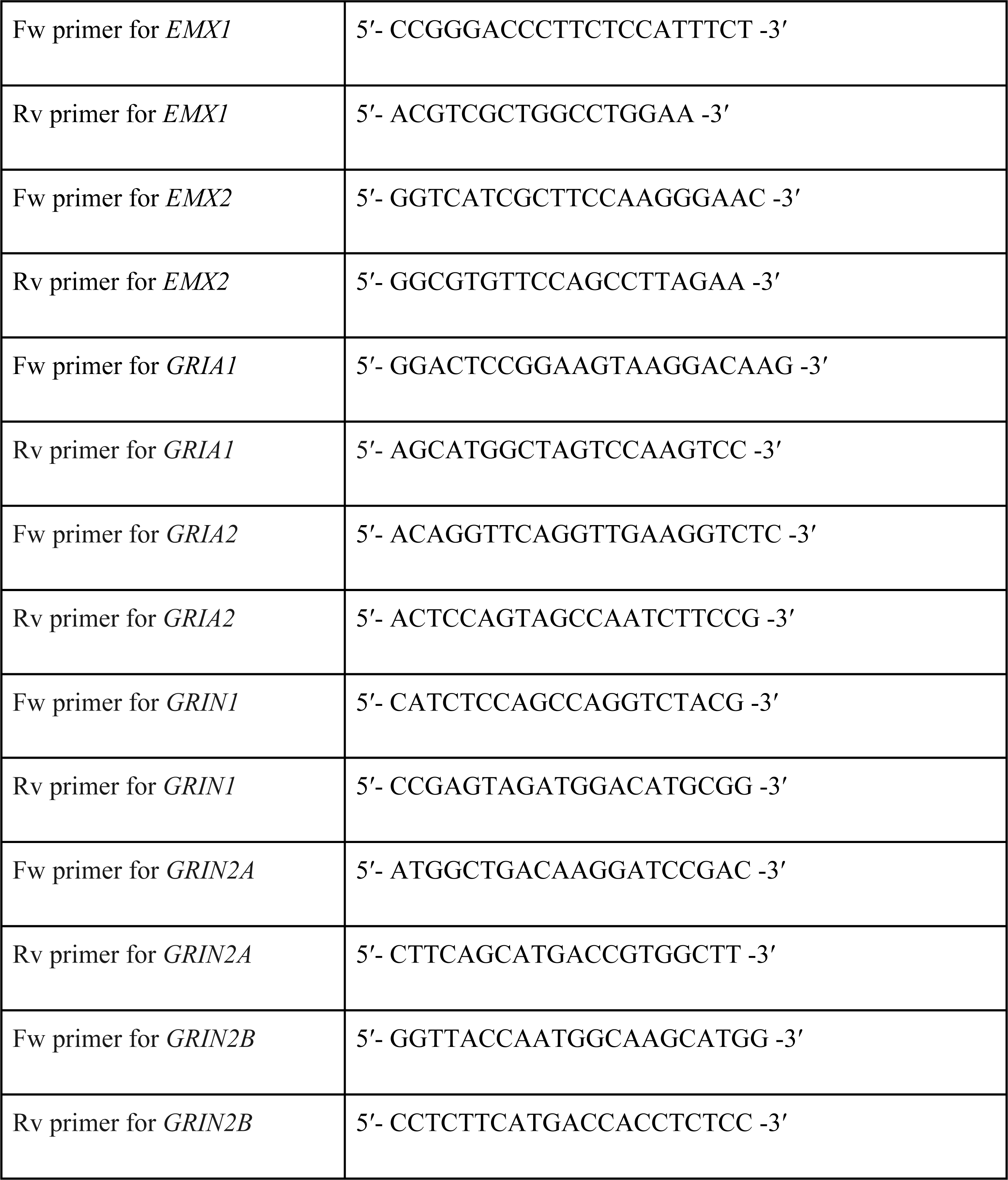

### Antibodies for immunohistochemistry

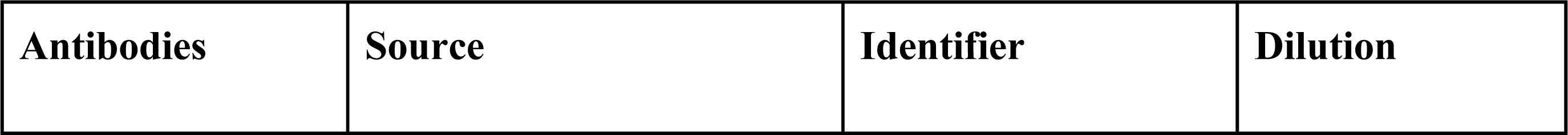
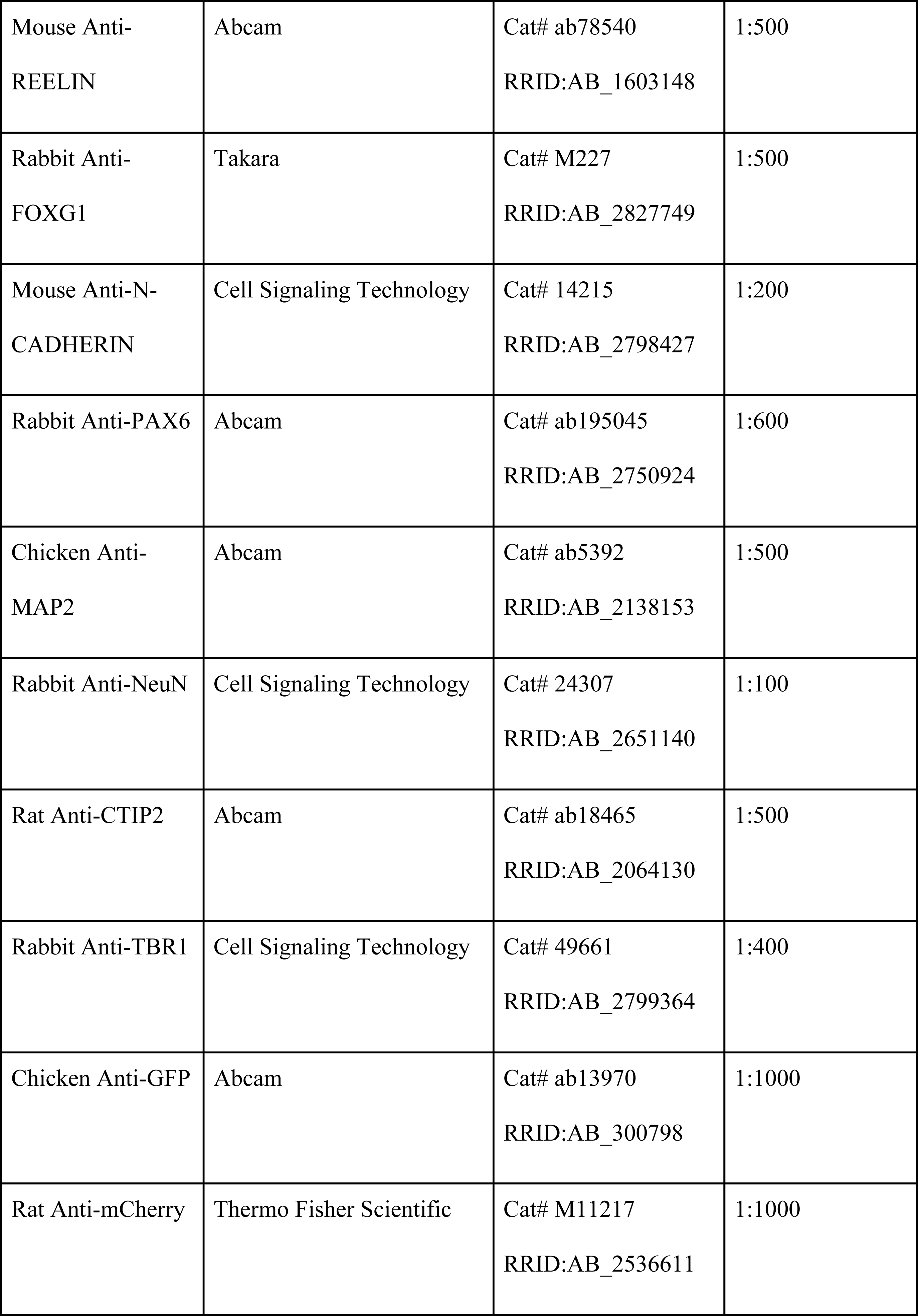
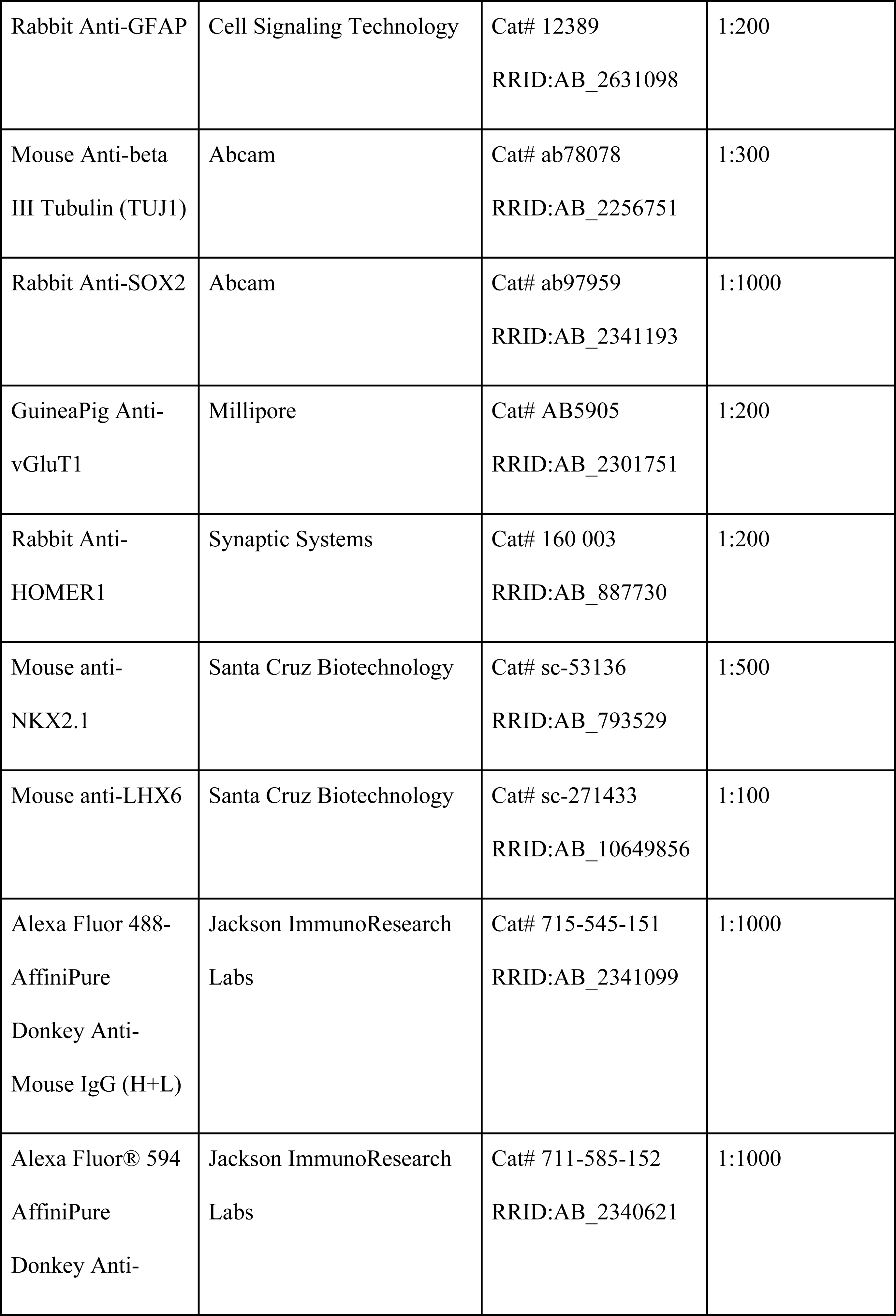
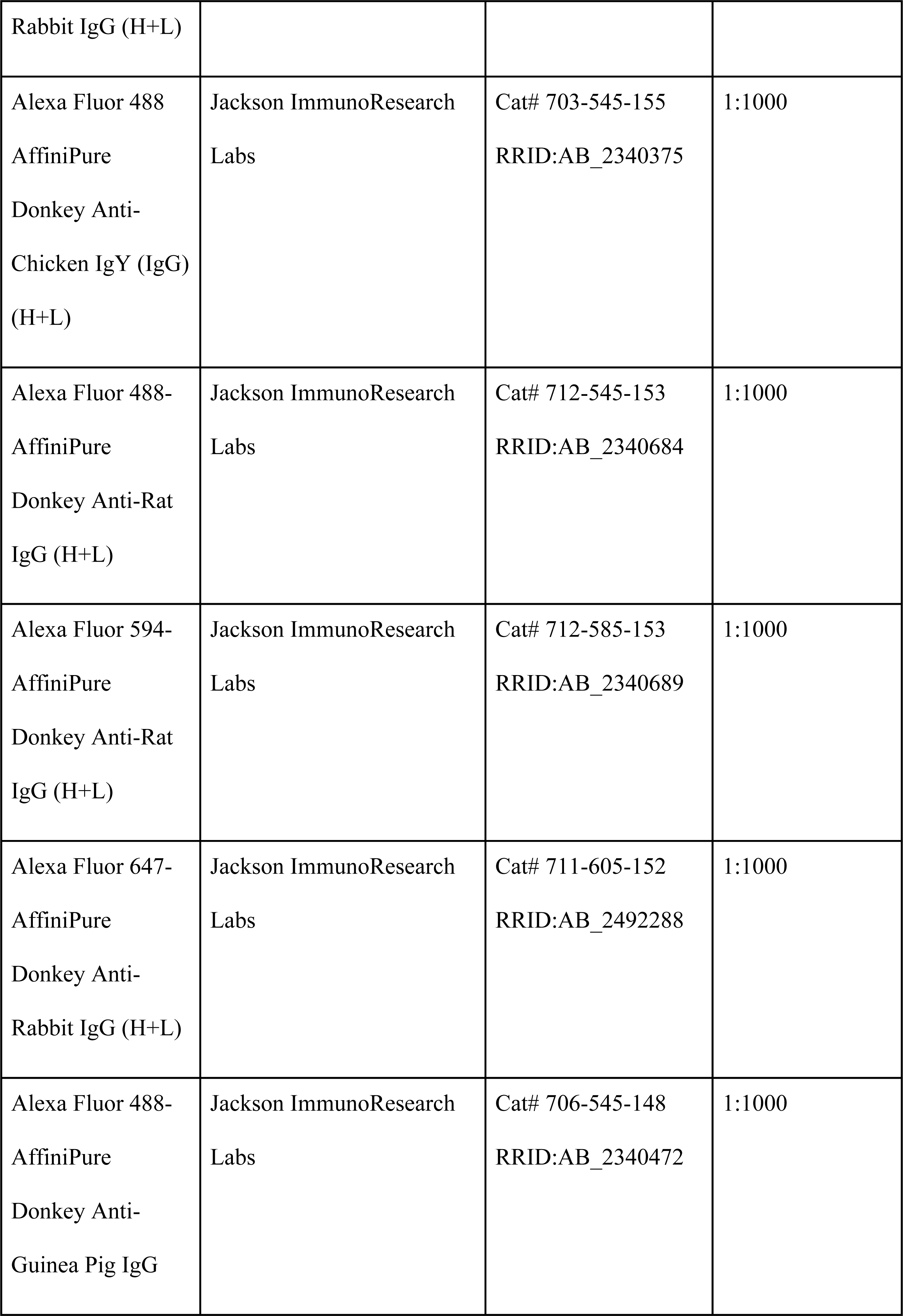
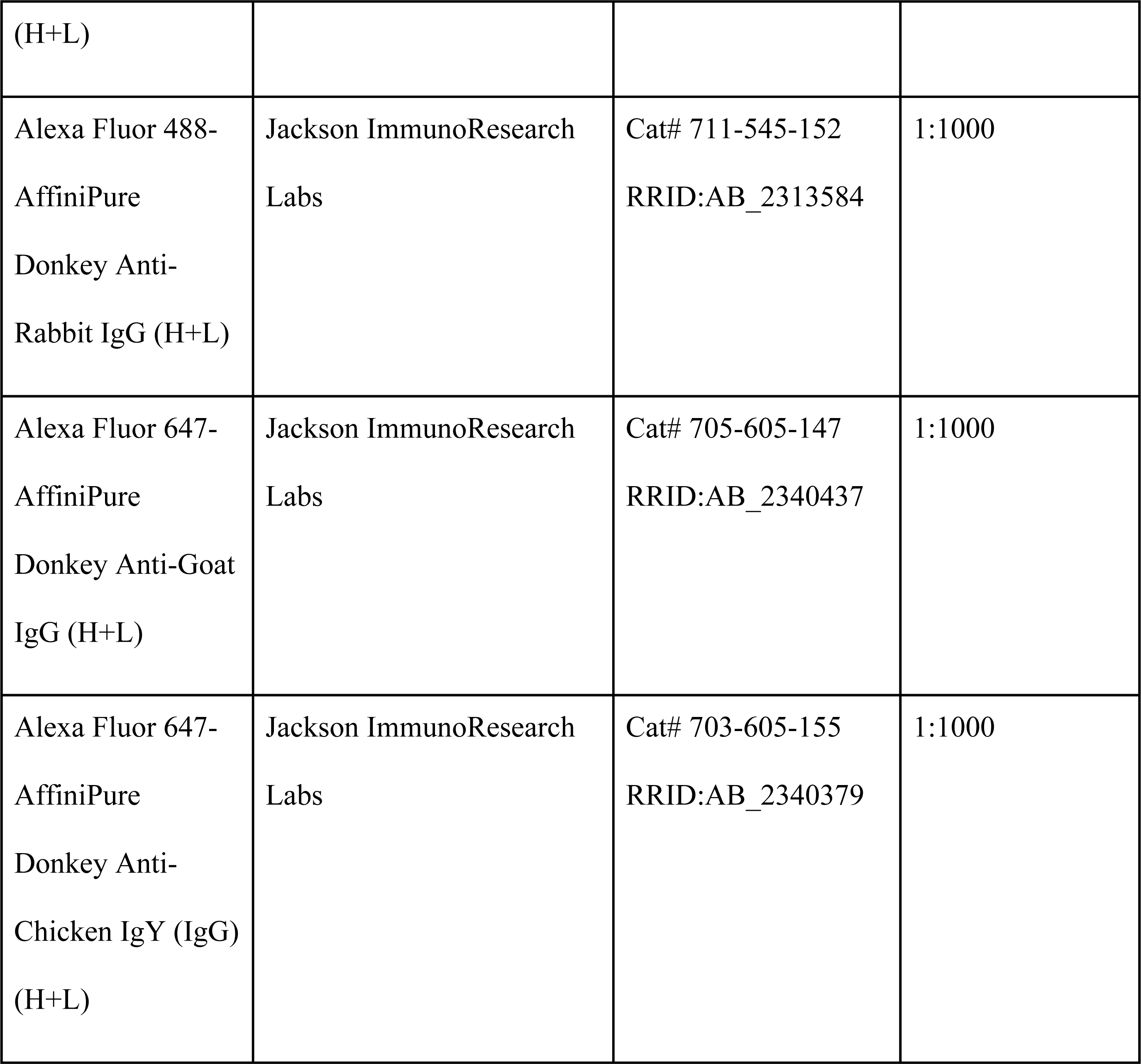

